# Defining the root endosphere and rhizosphere microbiomes from the World Olive Germplasm Collection

**DOI:** 10.1101/636530

**Authors:** Antonio J. Fernández-González, Pablo J. Villadas, Carmen Gómez-Lama Cabanás, Antonio Valverde-Corredor, Angjelina Belaj, Jesús Mercado-Blanco, Manuel Fernández-López

## Abstract

Up to date, the bacterial and fungal microbial communities from the olive (*Olea europaea* L.) root systems have not been simultaneously studied. In this work, we show that microbial communities from the olive root endosphere are less diverse than those from the rhizosphere. But more relevant was to unveil that olive belowground communities are mainly shaped by the genotype of the cultivar when growing under the same environmental, pedological and agronomic conditions. Furthermore, *Actinophytocola*, *Streptomyces* and *Pseudonocardia* are the most abundant bacterial genera in the olive root endosphere, *Actinophytocola* being the most prevalent genus by far. In contrast, *Gp6*, *Gp4*, *Rhizobium* and *Sphingomonas* are the main genera in the olive rhizosphere. *Canalisporium*, *Aspergillus*, *Minimelanolocus* and *Macrophomina* are the main fungal genera present in the olive root system. Interestingly enough, a high proportion of so far unclassified fungal sequences at class level were detected in the rhizosphere. From the belowground microbial profiles here reported, it can be concluded that the genus *Actinophytocola* may play an important role in olive adaptation to environmental stresses. Moreover, the huge unknown fungal diversity suggests that there are still some fungi with important ecological and biotechnological implications that have yet to be discovered.

## 1. Introduction

The cultivated olive (*Olea europaea* L. subsp. *europaea* var. *europaea*) is not only one of the oldest domesticated trees^1^, but also constitutes one of the most important and outstanding agro-ecosystems in the Mediterranean Basin, shaped along millennia^2^. In this area, there is an olive belt with more than 10 million ha belonging to the countries of the coastal regions, which account for around 80% of the world’s total olive cultivation area^3^. In some of these countries such as Spain, the world’s largest olive oil and table olive producer, this woody crop has undisputable social, economic and agro-ecological relevance, accounting for almost 25% of the total olive trees and more than 37% of the world’s olive oil production^4^. In addition to its ecological and social importance, the main product obtained from this iconic tree (i.e. the virgin olive oil), has a number of health and nutritional benefits so that its consumption is increasing worldwide^5^.

Olive cultivation is threatened by several abiotic (e.g. soil erosion) and biotic (i.e. attacks from insects, nematodes and pathogenic microbes) constraints. Among relevant phytopathogens present in the soil microbiota affecting olive health, representatives of the *Oomycota* class (e.g. *Phytophthora* spp.) as well as higher fungi (e.g. *Verticillium dahliae* Kleb.) must be highlighted^2,6–8^. In addition to the traditional and well-known microbiological menaces affecting olive crop (e.g., anthracnose [caused by *Colletotrichum* spp.], Verticillium wilt [VWO, *V. dahliae*], peacock spot [*Spilocea oleagina* (Cast.) Hughes.], or knot disease [*Pseudomonas savastanoi* pv. savastanoi Smith.])^9–12^, emerging diseases like the olive quick decline syndrome caused by *Xylella fastidiosa* Wells. ssp. *pauca* observed for the first time attacking olive trees in Italy in 2013^13^, must be considered. In addition to new threats, some reports warn on the increase in pathogen and arthropod attacks as a consequence of changing from traditional olive cropping systems to high-density tree orchards. However, the impacts of high-density olive groves on, for instance, soil-borne diseases have not been yet studied^14^. Another important menace to take into account is climate change, which is expected to affect the incidence and severity of olive diseases^6^. Finally, the reduction in the number of planted olive cultivars due to either commercial (e.g. improved yield, etc.) or phytopathological (i.e. tolerance to diseases) reasons, a trend observed in many areas, will eventually lessen olive genetic diversity. All these factors may have a profound, yet not evaluated impact on the composition, structure and functioning of belowground microbial communities^8^.

A comprehensive knowledge of microbial communities associated to the olive root system, including the root endosphere and the rhizosphere soil, is therefore instrumental to better understand their influence on the development, health and fitness of this tree. A priori, the vast majority of the olive-associated microbiota must be composed of microorganisms providing either neutral or positive effects to the host. Indeed, recent literature provides solid evidence that olive roots are a good reservoir of beneficial microorganisms, including effective biocontrol agents (BCA)^15–18^. Among the beneficial components of the plant-associated microbiota, endophytic bacteria and fungi are of particular interest to develop novel biotechnological tools aiming to enhance plant growth promotion and/or control of plant diseases. Moreover, microorganisms able to colonize and endure within the plant tissue pose the additional advantage to be adapted to the specific microhabitat/niche where they can provide their beneficial effects^19^. Besides endophytes, beneficial components of tree root-associated microbiota colonizing the rhizoplane and/or the rhizosphere soil can also directly promote plant growth (i.e. bio-fertilization, phyto-stimulation) or alleviate stress caused by either abiotic (i.e.environmental pollutants, drought, salinity resistance) or biotic (see above) constraints^8^.

Our knowledge on olive-associated microbiota is still very scarce and fragmentary. So far, bacterial communities associated with wild olive (*Olea europaea* L. subsp. *europaea* var. *sylvestris*) roots (endo- and rhizosphere) have been studied using fluorescent terminal restriction fragment length polymorphism (FT-RFLP) as culture-independent approach as well as bacteria isolation in culturing media^15^. In another study, endophytic fungi from the phyllosphere and roots of olive cultivar (cv.) Cobrançosa^20^ were screened by a culture-dependent method to compare the fungal communities between above- and belowground compartments. Microbial communities of the olive phyllosphere and carposphere have been analyzed using denaturing gradient gel electrophoresis (DGGE)^21^, isolation of fungi in culturing media^22^ and high-throughput sequencing of both fungal^23^ and prokaryotic^24^ communities.

In this study we aim, for the first time, to unravel the composition and structure of belowground prokaryotic and fungal communities of cultivated olive by high-throughput sequencing. A core collection of olive cultivars (36 originating from 9 different countries, Table 1) present at the World Olive Germplasm Collection (WOGC; Córdoba, Spain) representative of enough genetic diversity within the Mediterranean Basin have been analyzed when all varieties were grown under the same climatic, pedological and agronomic conditions. The following objectives were pursued: a) to perform an in-depth study of the belowground microbial communities (root endosphere and rhizosphere) in a wide range of olive genotypes; b) to assess what is/are the determinant factor(s) contributing to build up such communities; c) to establish the core and accessory microbiota of the olive rhizosphere and root endosphere. The hypothesis to-be-tested is that under specific agro-climatic and edaphic conditions the olive genotype is the key factor for building up the root endosphere community but not that important in the rhizosphere community.

**Table 1.**
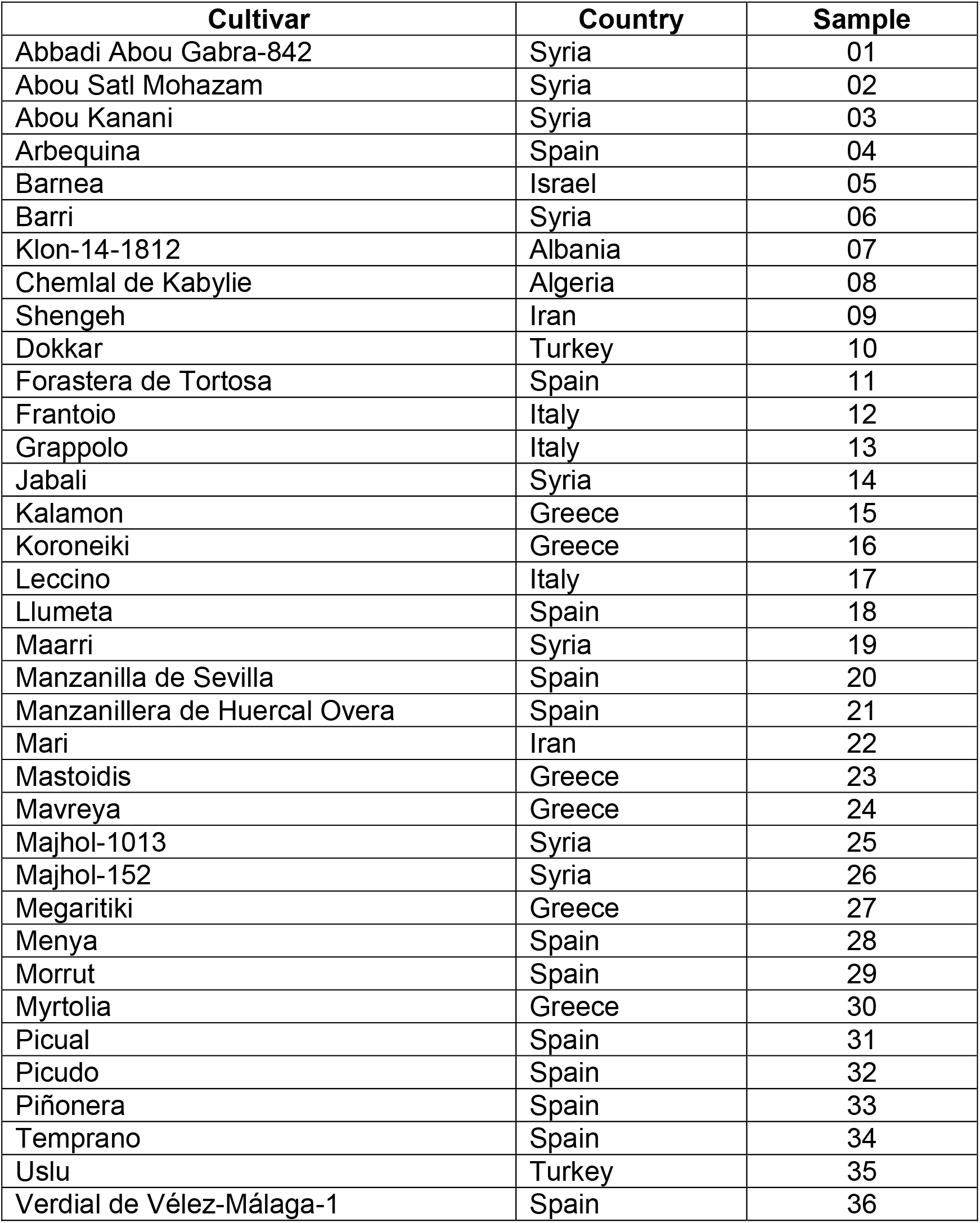
The 36 olive cultivars sampled in the World Olive Germplasm Collection (WOGC).

## 2. Results

### 2.1. Microbial communities clustered by compartments (endosphere and rhizosphere), and by olive cultivar in each compartment

Starting from about 37 million reads, a total of 1,404,769 (prokaryotic) and 1,005,148 (fungal), and 5,330,385 (prokaryotic) and 912,302 (fungal) good quality reads from the root endosphere and the rhizosphere, respectively, were eventually retained from the 36 olive cultivars here analyzed (Table 1). The smallest samples had 2,061 prokaryotic and 442 fungal sequences, coming from the root endosphere, and the largest ones reached 78,913 prokaryotic and 55,072 fungal sequences, these originated from the rhizosphere (Tables S1 and S2). After rarefying to the smallest sample, alpha diversity indices showed statistically significant differences between the two compartments (i.e. the endosphere and the rhizosphere), showing the rhizosphere samples the highest richness and diversity values (Figure S1). Subsequently, both compartments were split and rarefied independently for further alpha diversity analyses to 2,061 (442 in fungi) and 15,565 (665 in fungi) sequences from endosphere and rhizosphere, respectively.

With regards to the prokaryotic communities, richness showed significant differences when comparing the root endosphere of olive cultivars, showing just marginal differences in diversity. Considering the rhizosphere, only the diversity showed statistically significant differences among cultivars (Table 2, Figures S2a and b). Concerning the fungal communities, both richness and diversity indices showed statistically significant differences when comparing olive cultivars for each compartment (Table 2, Figures S2c and d).

**Table 2.** Comparisons of alpha diversity indices in the different microbial communities.

| **Prokaryotes** | Cultivar |  | Endosphere vs rhizosphere |
| --- | --- | --- | --- |
| Index | Root endosphere | Rhizosphere | Whole community |
| $S_{obs}$ | **0.0178 (36.8)** | 0.0500 (49.9) | **< 2.2e<sup>-16</sup> (122.2)** |
| Chao1 | **0.0357 (34.1)** | 0.2117 (41.4) | **< 2.2e<sup>-16</sup> (122.2)** |
| Shannon | 0.0774 (30.8) | **4.6e<sup>-05</sup> (77.6)** | **< 2.2e<sup>-16</sup> (122.2)** |
| InvSimpson | 0.0602 (31.9) | **8.5e<sup>-05</sup> (83.2)** | **< 2.2e<sup>-16</sup> (122.2)** |
| df | 21 | 35 | 1 |

| **Fungi** | Cultivar |  | Endosphere vs rhizosphere |
| --- | --- | --- | --- |
| Index | Root endosphere | Rhizosphere | Whole community |
| $S_{obs}$ | **0.0018 (60.4)** | **0.0096 (57.5)** | **< 2.2e<sup>-16</sup> (147.1)** |
| Chao1 | **0.0133 (52.3)** | **0.0119 (56.6)** | **< 2.2e<sup>-16</sup> (142.5)** |
| Shannon | **0.0014 (61.3)** | **0.0276 (52.8)** | **< 2.2e<sup>-16</sup> (110.9)** |
| InvSimpson | **0.0127 (52.5)** | 0.0593 (48.9) | **< 2.2e<sup>-16</sup> (82.8)** |
| df | 32 | 35 | 1 |

$S_{obs}$ : Observed richness
df: degree of freedom
In bold: significant p-values considering a confidence interval of 95%
In brackets: chi-squared values

We compared the distribution of the samples from both compartments, rhizosphere and root endosphere. Results showed significantly different prokaryotic (PERMANOVA R^2^ 0.43; p-value < 0.0001) and fungal (PERMANOVA R^2^ 0.06; p-value < 0.0001) communities (Figure 1a and b). Regarding to the prokaryotic root endosphere, the olive cultivar explained 42% of the variation, being statistically significant (PERMANOVA R^2^ 0.42; p-value < 0.0001) (Figure 2). Regarding to prokaryotic communities present in the olive rhizosphere, the cultivar explained more than 53% of the distribution, being statistically significant as well (PERMANOVA R^2^ 0.53; p-value < 0.0001) (Figure 3).

**Figure 1.**
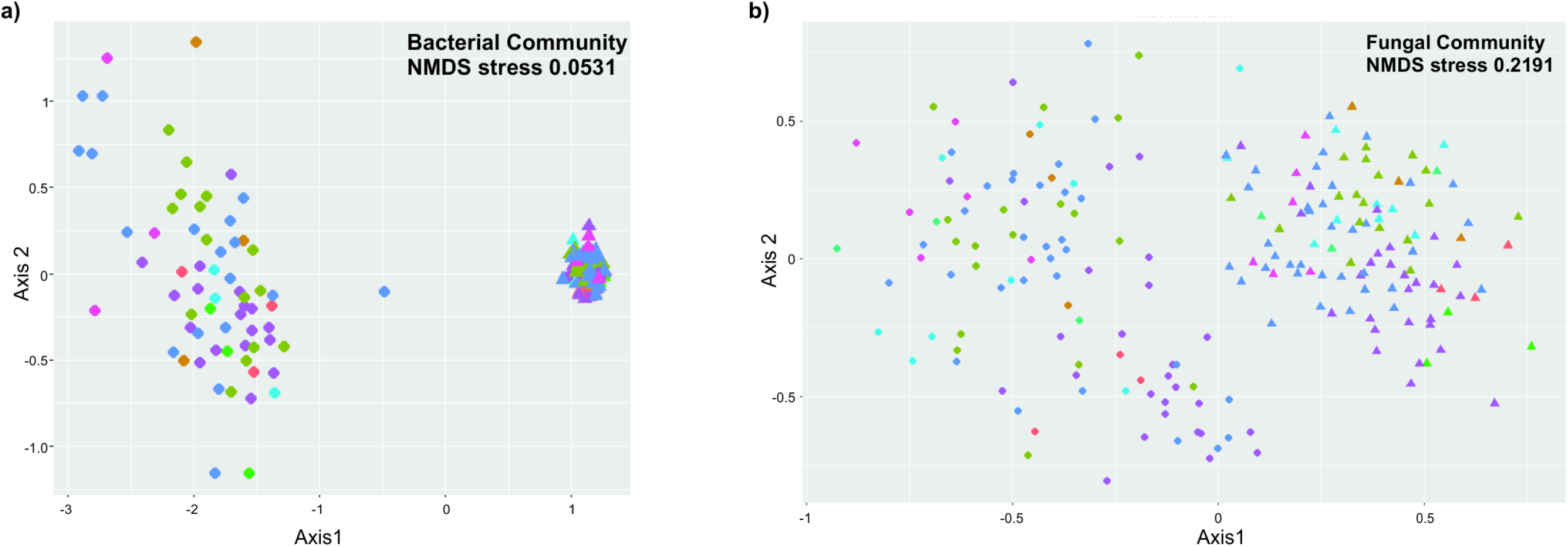
NMDS of bacterial (a) and fungal (b) communities by compartment. The letters A, B and c after the numbers were used to distinguish the 3 replicates of each cultivar. The different colors indicate the country of origin of the cultivars.

**Figure 2.**
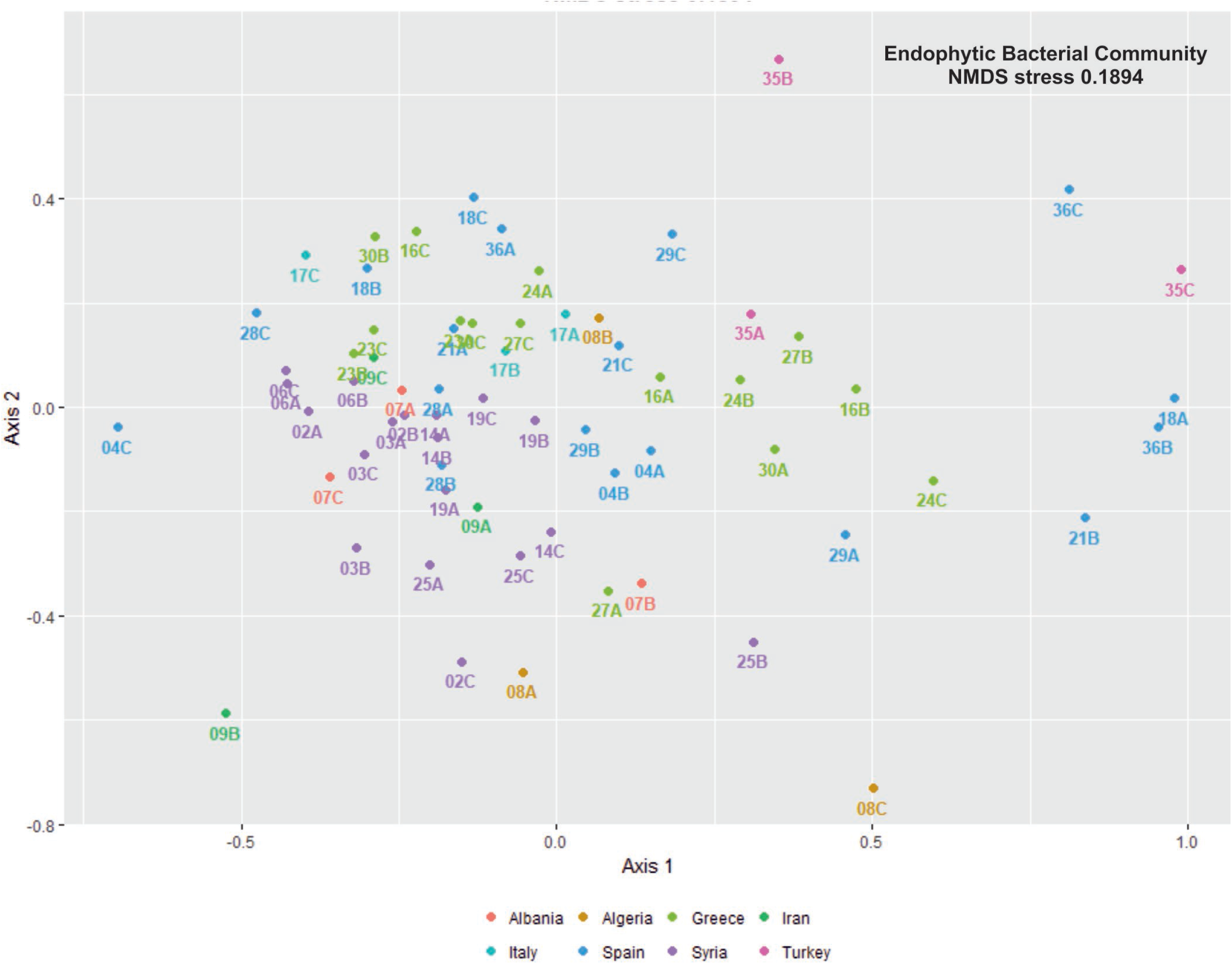
NMDS of bacterial communities from rhizosphere. The letters A, B and C after the numbers were used to distinguish the 3 replicates of each cultivar. The different colors indicate the country of origin of the cultivars.

**Figure 3.**
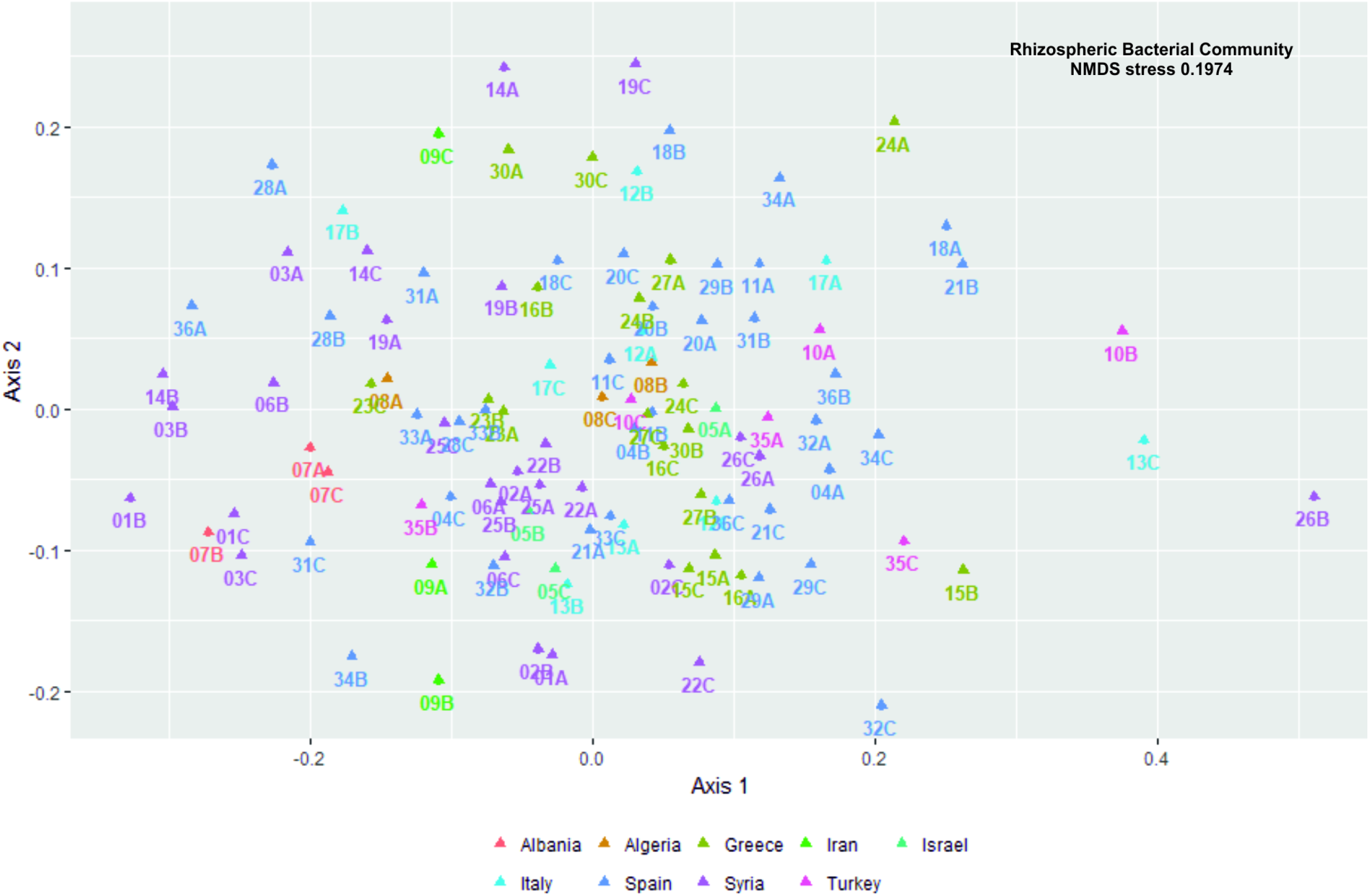
NMDS of bacterial communities from root endosphere. The letters A, B and C after the numbers were used to distinguish the 3 replicates of each cultivar. The different colors indicate the country of origin of the cultivars.

Concerning fungal communities in the root endosphere, the cultivar explained 39% of the variation, being statistically significant (PERMANOVA R^2^ 0.39; p-value < 0.0001). In the case of the fungal rhizosphere, this factor explained 44% of the variation, also being statistically significant (PERMANOVA R^2^ 0.44; p-value < 0.0001). Data not plotted because of the high NMDS stress value (0.22 with 3 dimensions).

### 2.2. The olive root endosphere and soil rhizosphere show different prokaryotic taxonomic profiles

Completely different taxonomic profiles at phylum level (class level for *Proteobacteria*) were obtained when comparing the prokaryotic communities residing in the olive root endosphere with those ones present on the olive rhizosphere (Figure 4). Despite the fact that universal primers for both prokaryotic kingdoms were used, no OTU was classified as *Archaea*. On the one hand, predominant phyla (or class) in the endophytic communities of the 22 olive cultivars examined (see Methods for exclusion criteria) were *Actinobacteria*, *Alphaproteobacteria*, *Gammaproteobacteria*, *Bacteroidetes* and *Deltaproteobacteria*, accounting for more than 90% of the sequences. Remarkably, *Actinobacteria* exceeded 50% in all of them, highlighting cultivars Chemlal de Kabylie, Llumeta and Mavreya (from Algeria, Spain and Greece, respectively) that represented more than 80% of the total number of sequences (Figure 4a). On the other hand, rhizospheric communities showed more even profiles with the phylum *Acidobacteria* accounting for an average of 27.5% of the sequences in the 36 olive cultivars here examined. *Acidobacteria* was followed by *Alphaproteobacteria* (18.8%), *Actinobacteria* (9.8%), *Gemmatimonadetes* (5.2%) and *Betaproteobacteria* (4.5%). Overall, and on average, the sum of all of them represented nearly 70% of the total number of sequences (Figure 4b). In contrast, *Gemmatimonadetes* and *Betaproteobacteria* were minor phyla in the olive root endosphere (0.06% and 0.8%, respectively).

**Figure 4.**
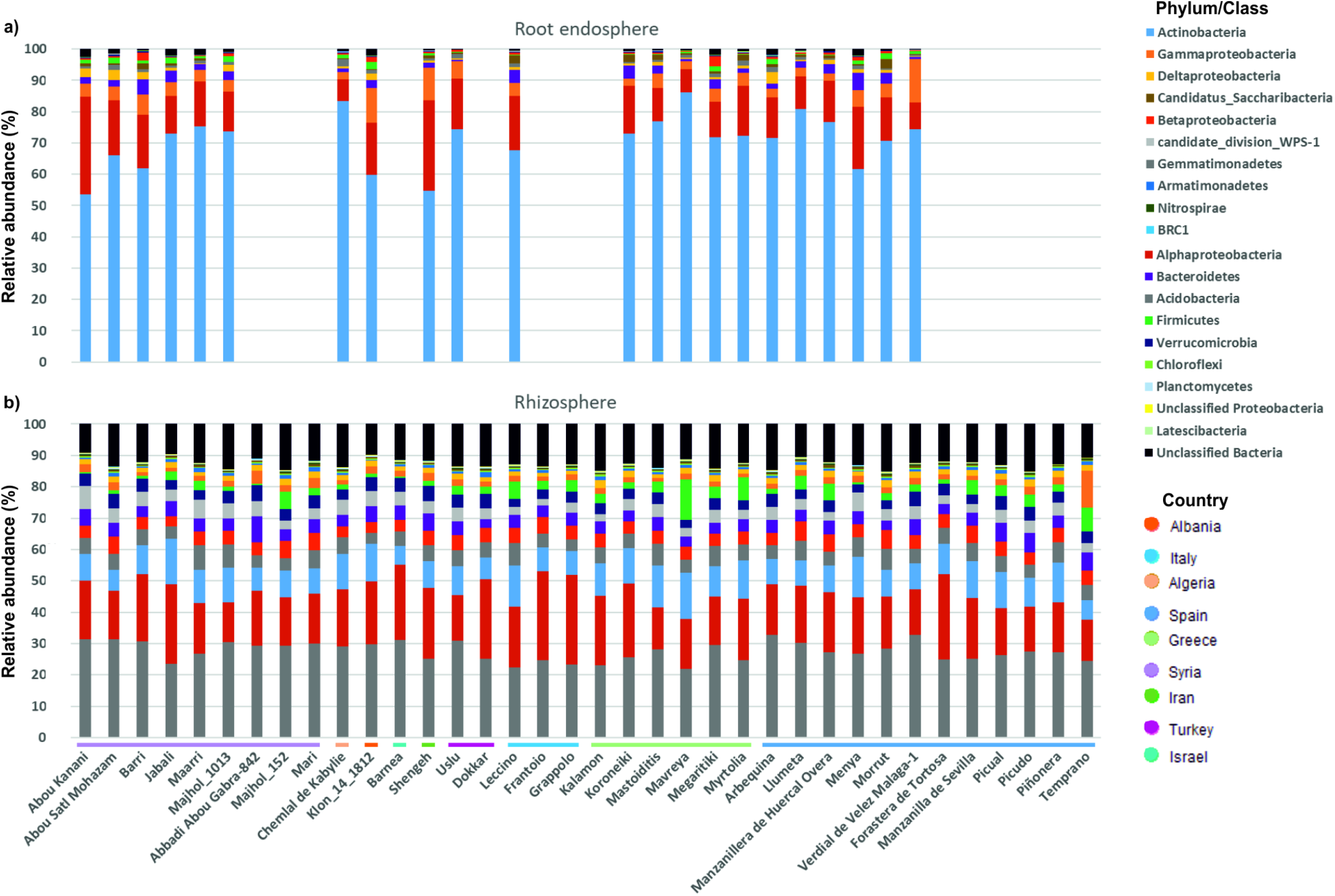
Bacterial phyla (class for *Proteobacteria*) in the root endosphere (a) and rhizosphere (b). The horizontal colored lines indicate the country of origin of the cultivars.

In the endosphere, and at the genus level, only two genera were significantly represented among the 22 cultivars that could eventually be examined: *Flavitalea* (*Bacteroidetes*) and *Actinophytocola* (*Actinobacteria*). Indeed, *Flavitalea* was most abundantly represented in cv. Myrtolia but absent in cv. Uslu (Figure S3a). Conversely, *Actinophytocola* was highly prevalent in 8 cultivars including Uslu (the highest) and Myrtolia (Figure S3b). Furthermore, *Actinophytocola* was the most abundant genus inhabiting the olive root endosphere accounting for an average of 22.1 ± 15.0% of the sequences, followed by *Streptomyces* (13.2 ± 8.2%), *Pseudonocardia* (9.4 ± 3.8%), *Bradyrhizobium* (2.6 ± 1.4%), *Ensifer* (2.6 ± 6.6%) and *Rhizobium* (2.0 ± 2.8%). The sum of relative abundances of these six main endophytic genera ranged from 33.3% in cv. Barri (Syria) to 73.1% in cv. Uslu (Turkey) (Figure S3c).

With regards to rhizosphere soil samples, our results showed that 63 genera were significantly more abundant among the different olive cultivars examined (36 cultivars). Moreover, eight out of the eleven main rhizosphere genera, with a relative abundance higher than 1%, had statistically significant differences between cultivars (Figure S4). Three of the most prevalent genera, namely *Gp6*, *Gp4* and *Gp7*, belong to the main rhizospheric phylum *Acidobacteria*, but only the first two showed significant differences. Belonging to the second most abundant phylum, *Proteobacteria*, and being both *α- Proteobacteria*, *Rhizobium* and *Sphingomonas* were also relatively highly abundant and showed significant differences among cultivars. The relative abundance of these eleven genera ranged from 49.4% in cv. Temprano (Spain) to 64% in cv. Barnea (Israel) (Figure S4).

### 2.3. Fungal taxonomic profiles only showed minor differences between the olive root endosphere and the soil rhizosphere

In contrast to prokaryotic communities, fungal communities showed more similar taxonomic profiles at the class level. The main difference between the two compartments was the percentage of sequences that remained unclassified (10.7% in the root endosphere *versus* 35.4% in the rhizosphere) (Figure 5). This proportion was very heterogeneous among olive cultivars, Grappolo (Italy) and Chemlal de Kabylie (Algeria) being the two cultivars that harbored more unclassified sequences in the root endosphere (37.8 and 29.4%, respectively), while cultivars Shengeh (Iran) and Abou Kanani (Syria) did so in rhizosphere samples (87.3 and 82.8%, respectively). The prevalent classes present in the olive root endosphere were *Sordariomycetes* (38.1%), *Eurotiomycetes* (23%), *Agaricomycetes* (13.2%) and *Dothideomycetes* (11.5%), accounting for more than 85% of the sequences obtained from the 33 olive cultivars assessed (see Methods for exclusion criteria) (Figure 5a). The remaining classes were clearly less relatively abundant, *Glomeromycetes* being the only one reaching 1%, on average, in all cultivars. Nevertheless, due to the heterogeneity found among the cultivars, this class represented more than 12 and 8% of relative abundance in the Syrian cvs. Maarri and Jabali, respectively.

**Figure 5.**
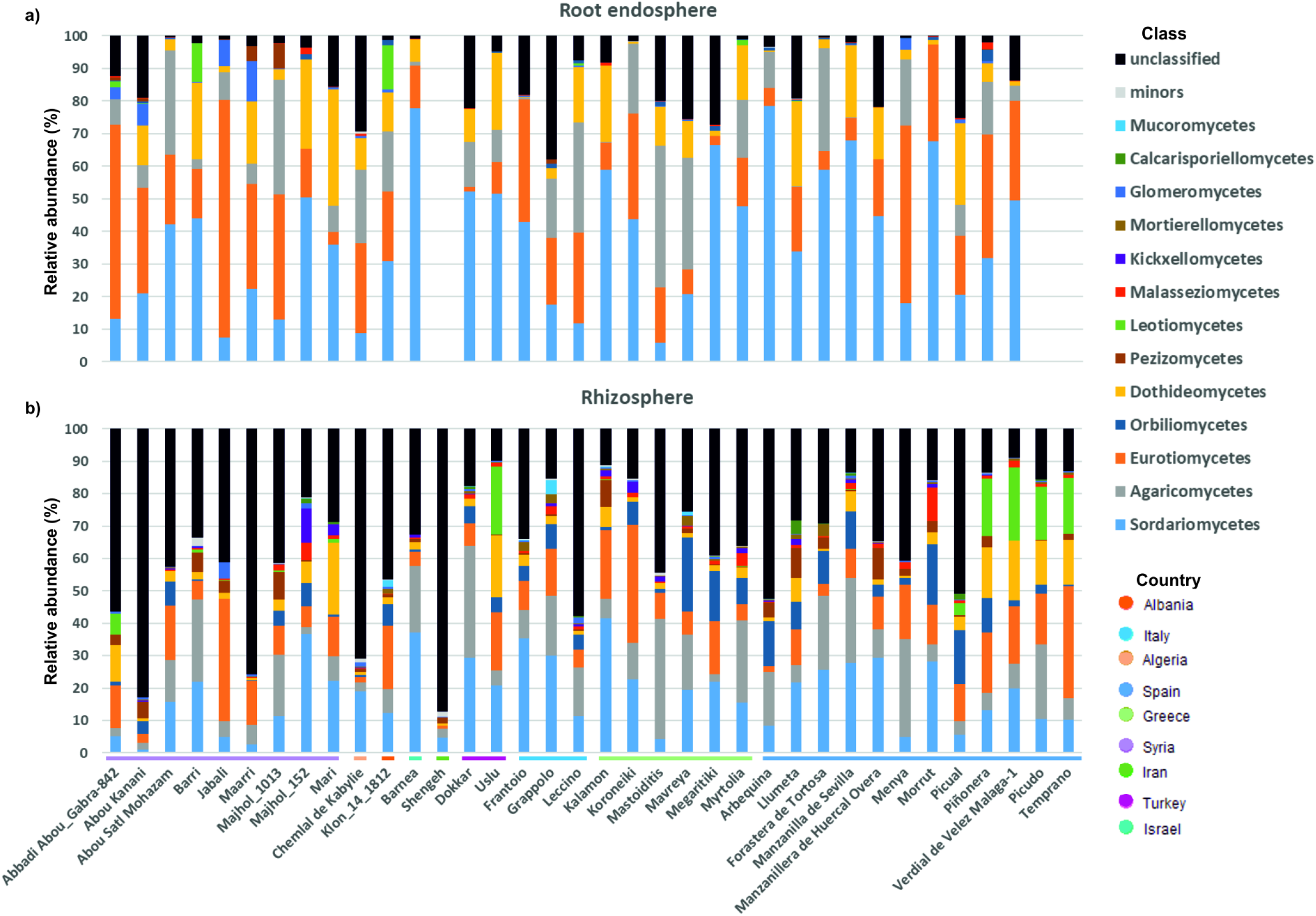
Fungal class in the root endosphere (a) and rhizosphere (b). The horizontal colored lines indicate the country of origin of the cultivars.

Regarding to rhizosphere communities, a smaller difference between the prevalent classes and the remaining ones was found in comparison to those found in the root endosphere. Similar to endophytic communities, *Sordariomycetes* was the predominant class in the rhizosphere (19%), followed by *Agaricomycetes* (12.9%), *Eurotiomycetes* (12.2%), *Orbiliomycetes* (6.5%) and *Dothideomycetes* (4.9%). While *Pezizomycetes* (2.4%) was, overall, more abundant than *Leotiomycetes* (2.3%) this latter class was exceptionally more abundant in the Spanish cultivars Piñonera, Picudo, Verdial de Velez Málaga-1 and Temprano (relative abundance ranging from 16.5 to 22.6%), and in the Turkish cv. Uslu (21.1%) (Figure 5b).

Concerning the genus level, only five fungal genera with statistically significant differences in relative abundance, *Scutellinia* (*Pezizomycetes*), *Acaulium*, *Purpureocillium* (*Sordariomycetes*), *Entoloma* (*Agaricomycetes*) and *Minimelanolocus* (*Eurotiomycetes*) were found in the root endosphere of the 33 olive cultivars examined (Figure S5a). Ten fungal endophytic genera were found with relative abundance higher than 1%, accounting for an average proportion of sequences ranging from 24.7% in cv. Grappolo (Italy) to 97.4% in cv. Forastera de Tortosa (Spain) (Figure S5c). However, due to the high heterogeneity of relative abundances observed for these genera, *Minimelanolocus* was the only genus showing statistically significant differences.

We found 7 fungal genera with statistically significant differences in relative abundance, *Macrophomina*, *Polyschema* (*Dothideomycetes*), *Minimelanolocus*, *Spiromastix* (*Eurotiomycetes*), *Cunninghamella* (*Mucoromycetes*), *Chlorophyllum* (*Agaricomycetes*), and *Dichotomopilus* (*Sordariomycetes*), in the rhizosphere of the 36 olive cultivars examined (Figure S5b). Only *Macrophomina* and *Minimelanolocus* showed enough relative abundance to be considered as part of the main fungal rhizosphere genera. Furthermore, *Macrophomina* was the third most abundant, on average, in all olive cultivars, particularly in 5 cultivars from Spain (Picual, Piñonera, Verdial de Velez Málaga-1, Picudo and Temprano) and in one from Turkey (Uslu) (Figure S5d).

### 2.4. Defining the belowground core microbiota of olive trees

Regarding bacterial communities, 46 (root endosphere) and 109 (rhizosphere) genera were found in all examined cultivars of each compartment. Furthermore, 40 of them were found in all the cultivars and in both compartments. Interestingly, 26 genera had a relative abundance higher than 1% in at least one compartment. The top 10 genera in the core olive root bacterial microbiota (bacteriota) were *Actinophytocola*, *Streptomyces*, *Gp6*, *Gp4*, *Pseudonocardia*, *Rhizobium*, *Sphingomonas*, *Gemmatimonas*, *candidate division WPS-1* and *Gp7*, accounting for almost 50% of the sequences in each compartment (Table 3; Table S3). Finally, all the main bacterial genera found in both compartments (Figures S3c and S4) were part of the core olive belowground bacteriota.

**Table 3.** Main (relative abundance ≥ 1%) core bacterial and fungal genera.

| Bacterial core |  |  |
| --- | --- | --- |
| Genus | Root endosphere (%)<sup>1</sup> | Rhizosphere (%)<sup>2</sup> |
| *Actinophytocola* | 22.07 | 0.07 |
| *Streptomyces* | 13.17 | 0.31 |
| *Gp6* | 0.58 | 11.08 |
| *Gp4* | 0.26 | 9.31 |
| *Pseudonocardia* | 9.37 | 0.14 |
| *Rhizobium* | 2.00 | 7.71 |
| *Sphingomonas* | 0.77 | 5.92 |
| *Gemmatimonas* | 0.06 | 5.24 |
| *candidate_division_WPS-1*<sup>4</sup> | 0.08 | 3.92 |
| *Gp7* | 0.04 | 4.08 |
| *Bacillus* | 0.68 | 2.31 |
| *Bradyrhizobium* | 2.57 | 0.20 |
| *Ensifer* | 2.56 | 0.15 |
| *Rubrobacter* | 0.05 | 2.48 |
| *Subdivision3*<sup>4</sup> | 0.02 | 2.35 |
| *Steroidobacter* | 1.78 | 0.34 |
| *Candidatus_Saccharibacteria*<sup>4</sup> | 1.03 | 0.40 |
| *Saccharothrix* | 1.18 | 0.07 |
| *Ohtaekwangia* | 0.21 | 1.03 |
| *Mycobacterium* | 0.98 | 0.09 |
| *Nonomuraea* | 1.04 | 0.04 |
| Fungal core |  |  |
| Genus | Root endosphere (%)<sup>3</sup> | Rhizosphere (%)<sup>2</sup> |
| *Canalisporium* | 29.53 | 6.05 |
| *Macrophomina* | 10.93 | 2.44 |
| *Aspergillus* | 1.66 | 3.84 |
| *Malassezia* | 0.28 | 1.37 |

<sup>1</sup> Relative abundance average of 22 cultivars920 <sup>2</sup> Relative abundance average of 36 cultivars921 <sup>3</sup> Relative abundance average of 33 cultivars922 <sup>4</sup> Name of *phylum*/class to which this *incertae sedis* genus belongs

Regarding fungal communities, only 4 (root endosphere) and 8 (rhizosphere) genera were found in all examined cultivars. Interestingly, the 4 core endophytic fungal genera were also part of the rhizosphere fungal core. Only 5 genera had a relative abundance > 1% in at least one compartment. The 4 fungal genera constituting the olive belowground fungal core were *Canalisporium*, *Macrophomina*, *Aspergillus* and *Malassezia*. They represented more than 40% of the endophytic sequences, but only 13.08% of the rhizosphere sequences. Furthermore, the 8 core rhizosphere fungal genera represented only 15.88% of the sequences (Table 3; Table S4).

## 3. Discussion

In addition to the higher alpha diversity (richness and evenness) found in the olive rhizosphere microbial communities compared to that observed for the microbiota inhabiting the root endosphere, and the finding that quite different communities were found in each compartment, a common scenario described in several studies^25,26^, the following major results must be highlighted from our work. Concerning the endosphere, cultivars originating from Syria showed the highest diversity level in contrast to the Turkish cultivars that showed the lowest one. With regard to the rhizosphere, fungal communities associated to cultivars from Albania and Syria appeared as the most diverse, while the Iranian and Israeli cultivars harbored the least diverse communities. Rhizosphere bacterial communities were not different in richness but showed dissimilar evenness. As observed for fungal communities, cultivars from Iran and Israel were also the least diverse in their rhizosphere bacterial assemblages.

Results here presented are in overall agreement with the major conclusion reported by Müller et al.^24^, even though these authors focused on aerial organs. Indeed, they concluded that the structure of endophytic prokaryotic communities residing in aboveground tissues was mainly driven by the geographical origin of the olive cultivars evaluated (Eastern: Greece, Syria; Central: France, Italy, Tunisia; and Western Mediterranean: Portugal, Spain, Morocco). The same overall conclusion is inferred from our study, emphasizing that is based on a larger number of cultivars and from a wider geographical origin (36 cultivars from 9 different countries *versus* 10 olive cultivars and 9 wild olive trees from 8 different countries in the study of Müller and co-workers). However, the main factor in our study was the genotype (cultivar) of each sample. It is true that the geographical origin was a statistically significant factor too but its variation was nested in cultivar variation (data from PERMANOVA test). In addition, more detailed information was obtained in our study. Thus, communities harbored by olive cultivars originating from Greece (olive green; see colors and distribution in Figures 2 and 3) and Spain (blue) showed more similarities among them than to those from Syrian (purple) cultivars. Moreover, the Italian (light blue) cultivar was intermingled between these two clusters and the unique Turkish (pink) representative tested in our work appeared as distantly related to the Syrian genotypes. Although a distinction among different geographical origins was observed, these clusters did not correspond to a longitudinal gradient (eastern, central, western Mediterranean countries), as reported by Müller et al.^24^. Our results indicate that the endophytic and rhizosphere microbial (bacteria and fungi) communities are mainly shaped by the olive genotype. We therefore conclude that the genotype is the main factor shaping olive belowground microbial communities, this factor being more determinant for the rhizosphere than for the endosphere, and more crucial for the bacteriota than for the mycobiota (see PERMANOVA R^2^ in results).

*Proteobacteria* has been described as the predominant prokaryotic phylum (about 90% of the relative abundance) present in root endophytic communities^26,27^. The same was observed for prokaryotic communities of the olive phyllosphere^24^. However, in our study, *Proteobacteria* (26% average relative abundance) was clearly overcome by *Actinobacteria* (64% average relative abundance) in the root endosphere. A similar finding has also been reported in *Agave* spp., particularly during the dry season^25^. Interestingly, no sequences belonging to the kingdom *Archaea* were detected in the root endosphere in our study, in contrast to the results by Müller et al.^24^ who reported that *Archaea* was a major group in the olive phyllosphere. In this latter study as well as in ours, the reverse primer used was the same. However, the forward primer used in our study has 94.6% archaeal amplification efficiency^28^. *Archaea* representatives were not found in the olive rhizosphere indicating that, without excluding the potential bias introduced by the primer pair here used, this kingdom is poorly represented in the olive belowground microbiota at least at the sampling time and under environmental conditions in which olive trees are cultivated in the WOGC.

The olive-associated microbiome harbors an important reservoir of beneficial microorganisms that can be used as plant growth promotion and/or biocontrol tools^15,24^. Moreover, bacterial antagonists of olive phytopathogens isolated from the olive root endosphere or the rhizosphere have the advantage to be adapted to the ecological niche where they can potentially exert their beneficial effect^18^. For instance, *Proteobacteria* and *Firmicutes* representatives, usually found as natural inhabitants from the olive rhizosphere, are thus good examples of effective antagonists against *V. dahliae*^16–18,29^. Besides, representatives of these phyla such as the genera *Pseudomonas* and *Bacillus* are easy to isolate, manipulate, propagate and formulate as BCAs. In addition to these well-known genera, species of the actinobacterial genus *Strepytomyces* have also been demonstrated as excellent BCAs in different pathosystems^30,31^. Moreover, the potential biocontrol of non-streptomycete *Actinobacteria* genera has been reported as well^30,32–34^. Taking into account that the prevalence of *Actinobacteria* found in our study (the genera *Actinophytocola*, *Streptomyces* and *Pseudonocardia* ranged from 30 to 60% of the bacterial olive root endophytic community), the isolation and in-depth characterization of culturable representatives of these genera will be of interest for their assessment as potential PGPR and/or BCA against olive tree pathogens. The genus *Actinophytocola*, described for the first time in 2010^35^ as a root endophytic actinobacteria (*Pseudonocardiaceae* family), has been isolated from Saharan non-rhizosphere soils in the south of Algeria^34^. Interestingly enough, these authors demonstrated its antimicrobial ability against some bacteria and fungi. *Actinophytocola* sp., in addition to other actinomycetes, has also been demonstrated to inhibit the growth of well-known human pathogens (*B. subtilis* and *S. aureus*)^36^. Finally, *Actinophytocola gilvus* was recently isolated from extremely dry conditions, from a soil crusts sample collected in the Tengger Desert in China^37^. Considering that this genus was ubiquitously and abundantly found in our study, *Actinophytocola* spp. inhabiting olive roots can be relevant for olive fitness and health (i.e. drought tolerance, broad antimicrobial activity range, etc.), what grants further research efforts aiming to isolate and characterize members of this relevant component of the olive belowground microbiome.

This is the first study in which a high-throughput sequencing approach has been implemented to unravel the olive belowground fungal communities. *Sordariomycetes* (38%) and *Eurotiomycetes* (23%), both belonging to *Ascomycota*, were found as the most abundant fungal classes in the root endosphere of olive. *Sordariomycetes* was previously found as the main endophytic fungal class in olive roots using a culture-dependent approach^20^. The endophytic fungal communities earlier found in aboveground olive tree compartments (phyllosphere and carposphere) by high-throughput sequencing^23^ or a culture-dependent approach^20^, differed from belowground communities reported in our study. This outcome reinforces previous reports showing important differences between above- and belowground olive fungal communities, regardless the methodological approach implemented to study them^20,22,23^. Interestingly enough, *Sordariomycetes* was also the most abundant class found in olive fruits regardless or not the presence of anthracnose symptoms^23^, pointing to the fact that this fungal class seems to be ubiquitously colonizing the interior of olive tissues. In the rhizosphere, *Agaricomycetes* (12.7%), belonging to *Basidiomycota*, and *Eurotiomycetes* (12.6%) were the most abundant classes. At this taxonomic level, the main difference between the two compartments was the percentage of unclassified sequences (12.4% in root endosphere and 35.6% in rhizosphere). Furthermore, in the particular case of cvs. Abou Kanani and Shengeh, unclassified sequences represented more than 80% of the good quality sequences found in the rhizosphere. The high percentage of unclassified sequences in this compartment seems to be a common finding^38^, although less pronounced in annual plants^39^, when using the same fungal database. According to previous studies and data here obtained, the olive rhizosphere carries a huge fungal diversity yet to be discovered. We have to take into account that much of those unclassified sequences do not belong to unknown fungi but, they were not properly classified due to limitation in the method and the database currently available. Notwithstanding, this may have important ecological implications for the tree, and pose novel agro-biotechnological avenues to be explored.

At the genus level, the structure and composition of olive belowground fungal communities also showed important differences compared to previous reports. For instance, in the particular case of phytopathogenic fungi, *Phomopsis columnaris* (fungus species causing of twig dieback of *Vaccinium vitis-idaea* [lingonberry])^40^ and *Fusarium oxyporum* were found by Martins et al.^20^ as the most relative abundant species, although sampled trees did not show visible symptoms. These results are far apart from ours. In our study, the above-mentioned species and the genera to which they belong were absent. However, the pathogenic fungi *Macrophomina phaseolina* showed relevant relative abundance in several cultivars and for both compartments. *Macrophina phaseolina* is a well-known pathogen causing charcoal rot in important crops including olive^16,41–44^, and it has also been shown that olive leaves produce compounds able to reduce its pathogenic activity^45^. This finding raises the possibility that *M. phaseolina* could be a common component of the olive-associated microbiota, but may reside within olive tissues without causing visible symptoms until external factors and/or microbiota alterations (dysbiosis) trigger a pathogenic stage. With regard to relevant soil-borne olive pathogens, it is worth mentioning that neither sequences corresponding to *Verticillium* spp. and *Fusarium* spp. nor to the oomycetes *Phytophthora* spp. and *Phytium* spp. were found in our study, confirming the good phytosanitary status in the WOGC soil. Finally, and regarding beneficial fungi, representatives of the genus *Trichoderma* were found in the rhizosphere of all cultivars but Chemlal de Kabylie and Llumeta. Species of this genus have been successfully used as BCA against VW of olive^46,47^.

In the olive belowground (endophytic and rhizosphere) core bacteriota here reported, genera from which some species have been well characterized and described as BCA are present. For instance, *Streptomyces* was the second most abundant genus in the endosphere whereas *Bacillus* was the tenth more abundant in the rhizosphere. While *Pseudomonas* was part of the rhizosphere core bacteriota, it was not considered as constituent of the endophytic core because it was absent in the root endosphere of cv. Mavreya. Nevertheless, *Pseudomonas* was relatively much more abundant inside olive root tissues than in the rhizosphere. Regarding the core mycobiota, and as mentioned above, the most noticeable presence of a pathogenic fungus was that of *Macrophomina*, and to a lesser extent *Colletotrichum.* The reported core microbiota indicates that, under the conditions found in the WOGC, olive trees harbor an important reservoir of beneficial/neutral microbes, and that the presence of deleterious microorganisms is nearly anecdotal. This correlates with the good development and appearance of the trees in the examined orchard, showing no visible symptoms of biotic stresses. The role of native microbiota in protecting plants from soil-borne pathogens has been highlighted in previous studies^48^. Nonetheless, further studies have to be carried out in the presence of soil-borne pathogens, such as *Verticillium dahliae*, to study the community alterations and confirm the protective role of some of the core microorganisms described in the present study.

## 4. Materials and methods

### 4.1. Sample Collection

Soil and root samples were collected from the World Olive Germplasm Collection (WOGC) (37°51’38.11’’ N; 4°48’28.61’’ W; 102 m.a.s.l.) located at the *Instituto de Investigación y Formación Agraria y Pesquera* (IFAPA, Córdoba, Spain) in the spring of 2017, when the trees were in full bloom. The selected 36 olive cultivars (Table 1) surveyed are grown in the same orchard to avoid differences related to the physicochemical characteristics of the soil, water availability, agricultural management, weather conditions or any other influencing factor. The cultivars selected represent the subset of the working olive core collection from the WOGC^49^. Geographical origin and commercial interest of varieties were the main criteria to choose these cultivars for downstream studies. The upper layer (first 5 cm) of soil was removed and rhizosphere soil samples were collected (5 to 20-cm depth) following the main roots of each plant until finding non-suberified roots. Root samples were also collected from the same plant to assess the root endophytic communities. Three soil and root samples from different trees of each cultivar were collected (n = 108). Furthermore, 10 bulk soil samples (1 kg) were collected at 1-1.5 m trunk distance of randomly selected trees (among the ones chosen for soil/root sampling) to analyze a number of physicochemical parameters of the WOGC soil (Table S5). These spots were randomly scattered along the orchard. Bulk soil samples in plastic bags were then transferred to the Agri-Food Laboratory of the Andalusian Regional Government at Córdoba (Spain), where physiochemical analyses were performed using standardized procedures.

### 4.2. DNA extraction and Illumina sequencing

The soil DNA from each individual sample was obtained using the Power Soil DNA Isolation kit (MoBio, Laboratories Inc., CA), following the manufacturer’s recommendations within 24 hours of samples collection. The root DNA was obtained, after root surface sterilization and grinding, using ‘Illustra DNA extraction kit Phytopure’ (GE Healthcare, Little Chalfont, UK). To ensure that DNA originated from endophytic microorganisms, and that microorganisms attached to the rhizoplane were eliminated, a thorough root surface sterilization protocol was implemented. Firstly, 20 ml of NaCl 0.8 % were added to 50 ml screw cap polypropylene tubes containing each root sample. Tubes were then vigorously shaken in order to remove adhering soil particles. After discarding the supernatant, roots were washed five times with distilled water. Secondly, the following root surface sterilization protocol was implemented: 70% ethanol for 5 min, NaClO (3.7%) containing Tween 20 0.01 % for 3 min, and finally 3 rinses in sterile and distilled water. To confirm that the disinfection protocol was successful, aliquots (100 µl) of water from the final rinse were plated in NA (Nutrient Agar) and PDA (Potato Dextrose Agar) plates that were incubated at 28°C for 7 days. Then, plates were examined to confirm the absence of microbial growth. DNA yields and quality were checked both by electrophoresis in 0.8% (w/v) agarose gels stained with GelRed and visualized under UV light, and using a Qubit 3.0 fluorometer (Life Technologies, Grand Island, NY). The DNA was sequenced with Illumina MiSeq platform in a commercial sequencing service (The Institute of Parasitology and Biomedicine “López Neyra”, CSIC, Granada, Spain). In the first run, a prokaryotic library was constructed amplifying the hyper-variable regions V3-V4 of the 16S rRNA gene using the primer pair Pro341F and Pro805R according to Takahashi et al.^28^. In the second run, an eukaryotic library was constructed amplifying the ITS1 region using the primer pair ITS1FI2 and ITS2 according to Schmidt et al.^50^ and developed by White et al.^51^. Both runs were sequenced using a paired-end 2×300-bp (PE 300) strategy. These sequence data have been submitted to the NCBI Sequence Read Archive (SRA) under the BioProject number PRJNA498945.

### 4.3. Data quality screening and overlapping

Demultiplexed and Phi-X174-free reads were quality checked with FastQC v.0.11.5^52^ and end-trimmed with FASTX-Toolkit v.0.014^53^. All the 3’ end nucleotides were removed until the first position which reached an average quality value bigger than Q25. The paired reads were overlapped with fastq-join v.1.3.1^54^ requesting a minimum overlap of 40 bp and a maximum of 15 % of difference in the overlapping region. Both libraries were processed with the same bioinformatics tools but following different pathways detailed below.

### 4.4. Prokaryotic data processing

The overlapped reads from the prokaryotic (Bacteria and Archaea) library were initially classified with an 80% bootstrap cutoff to the Ribosomal Database Project (RDP-II) 16S rRNA reference database, training set v.16 MOTHUR-formatted^55^, with MOTHUR v.1.39.5^56^. This initial step was performed to remove reads belonging to mitochondria, chloroplast and not identified at kingdom level (unknown). Then, using the software SEED2 v.2.1.05^57^ the prokaryotic sequences were trimmed and clustered. Firstly, by trimming the specific primers; then, by removing sequences with ambiguities and shorter than 400 bp as well as reads with an average read quality lower than Q30. Secondly, chimeric reads were removed by VSEARCH “De Novo” v.2.4.3^58^ implemented in SEED2 and OTUs were clustered with the same tool at 97% similarity. Finally, the OTU table was saved and OTUs accounting for less than 0.005% of the total sequences were removed according to Bokulich et al.^59^ for further analyses. The most abundant OTU sequences were retrieved in SEED2 and classified as mentioned above. This classification was considered as the taxonomic information of each OTU.

### 4.5. Eukaryotic data processing

The eukaryotic library was directly quality-trimmed in SEED2 by the removal of sequences with ambiguities and an average read quality lower than Q30. There was not size exclusion and the primers were initially kept for the next step. Subsequently, ITSx v.1.0.11^60^ was performed but the result was discarded because of it was unable to properly recognize and remove the forward primer (ITS1FI2). Then, to accurately extract the ITS1 region, the high quality reads were aligned against the ribosomal operons of *Saccharomyces cerevisiae* S288c using Geneious R11^61^. As expected, the forward primer plus 4 nt matched the end of the 18S rRNA gene, and the reverse primer plus 30 nt matched the beginning of the 5.8S rRNA gene. Both intragenic ends were removed using SEED2 and chimeric sequences identified and discarded with VSEARCH “De Novo” implemented in SEED2. Then, the good quality sequences were distance-based greedy clustered at 97% similarity with VSEARCH algorithm implemented in MOTHUR. The most abundant OTU sequences were classified using the UNITE v.7.2 dynamic database^62^ with MOTHUR following the parameters recommended in the website and used by Findley et al.^63^. Finally, only OTUs with more than 0.005% of the sequences and assigned to kingdom Fungi were kept for further analyses. Furthermore, OTUs assigned to the phylum Oomycota were manually checked to examine the (possible) presence of the phytopathogenic genera.

### 4.6. Statistical analyses

Alpha diversity indices (observed and Chao1 richness; Shannon and inverse of Simpson diversity) were compared with Kruskal-Wallis test and p-values were FDR corrected by the Benjamini-Hochberg method using the R package a*gricolae*^64^. Concerning the beta diversity, a normalization of the filtered OTU sequence counts was performed using the “trimmed means of M” (TMM) method with the BioConductor package *edgeR*^65^. The normalized data were considered to perform Nonmetric MultiDimensional Scaling (NMDS) on Bray-Curtis dissimilarities to ordinate in two dimensions the variance of beta diversity between compartments (root endosphere and rhizosphere) and among cultivars in each compartment, in both kingdoms. Ordination analyses were performed using the R package *phyloseq*^66^. We analyzed compartment and olive cultivar effects on community dissimilarities with permutational analysis of variance (PERMANOVA) and permutational analysis of multivariate dispersions (BETADISPER) using the functions *adonis* and *betadisper* in the *vegan* package with 9,999 permutations^67^. Significant prokaryotic or fungal genera in olive cultivar were obtained with the following protocol: i) we tested for differential genus abundance using likelihood ratio tests (LRT) in the normalized data with the R package *edgeR*; ii) we tested for differential genus abundance using proportions in non-normalized counts with the STAMP v.2.1.3 software^68^, selecting default statistical comparisons for multiple groups and firstly considering both Benjamini-Hochberg FDR for multiple test correction and without FDR correction; iii) those genera significantly different in the two methods previously described were plotted and manually checked to generate the final selection. Most of the steps performed on R were carried out following the R script publicly donated by Hartman et al.^69^.

## Acknowledgments

This work was supported by grant AGL2016-75729-C2-1-R from the Spanish Ministerio de Economía, Industria y Competitividad/Agencia Estatal de Investigación, and co-financed by the European Regional Development Fund (ERDF).

We thank Dr. José F. Cobo-Díaz from the Université de Brest, Laboratoire Universitaire de Biodiversité et Ecologie Microbieene IBSAM, ESIAB, Plouzané, France for technical support in eukaryotic data analysis; and to PhD student Ana V. Lasa from Departamento de Microbiología del Suelo y Sistemas Simbióticos, Estación Experimental del Zaidín – CSIC for technical support in R scripts.

## Author Contributions

JMB and MFL conceived the study and performed its design. AJFG, JMB and MFL wrote the manuscript. AB maintains the crop system and helped with sampling. PJV, CGLC and AVC conducted the sampling and DNA extractions. AJFG performed the bioinformatics analysis and analyzed the data. All authors read and approved the final manuscript.

## Supplementary Information

Table S1. Bacterial alpha diversity indices of each sample in both compartments. (xlsx)

Table S2. Fungal alpha diversity indices of each sample in both compartments. (xlsx)

Table S3. Core bacterial communities in the endosphere, rhizosphere and both at genus level. (xlsx)

Table S4. Core fungal communities in the endosphere, rhizosphere and both at genus level. (xlsx)

Table S5. Physicochemical properties of the soil from the World Olive Germplasm Collection (Córdoba, Spain). (docx)

Figure S1. Normalized alpha diversity indices by compartment in the prokaryotic (a) and the fungal (b) communities. Endosphere (Endo), Rhizosphere (Rhizo) and Richness (Observed). (pptx)

Figure S2. Microbial (bacterial a and b; fungal c and d) normalized alpha diversity indices of each sample clustered by cultivars in both compartments (a and c endosphere; b and d rhizosphere). (pptx)

Figure S3. Statistically significant endophytic bacterial genera by cultivar (a, b) and the main bacterial genera in the endosphere (c). (pptx)

Figure S4. Main bacterial genera in the rhizosphere. Asteriscs indicate statistically significant differences in the relative abundance when comparing cultivars. (pptx)

Figure S5. Statistically significant fungal endophytic (a) and rhizosphere (b) genera by cultivar and the main fungal genera in the endosphere (c) and the rhizosphere (d). The main fungal genera were highlighted with an asterisc to indicate statistically significant differences in the relative abunance when comparing cultivars. (pptx)

