## Supplementary Material for "Defining the root endosphere and rhizosphere microbiomes from the World Olive Germplasm Collection"

Figure S1. Normalized alpha diversity indices by compartment in the prokaryotic (a) and the fungal (b) communities. Endosphere (Endo), Rhizosphere (Rhizo) and Richness (Observed).

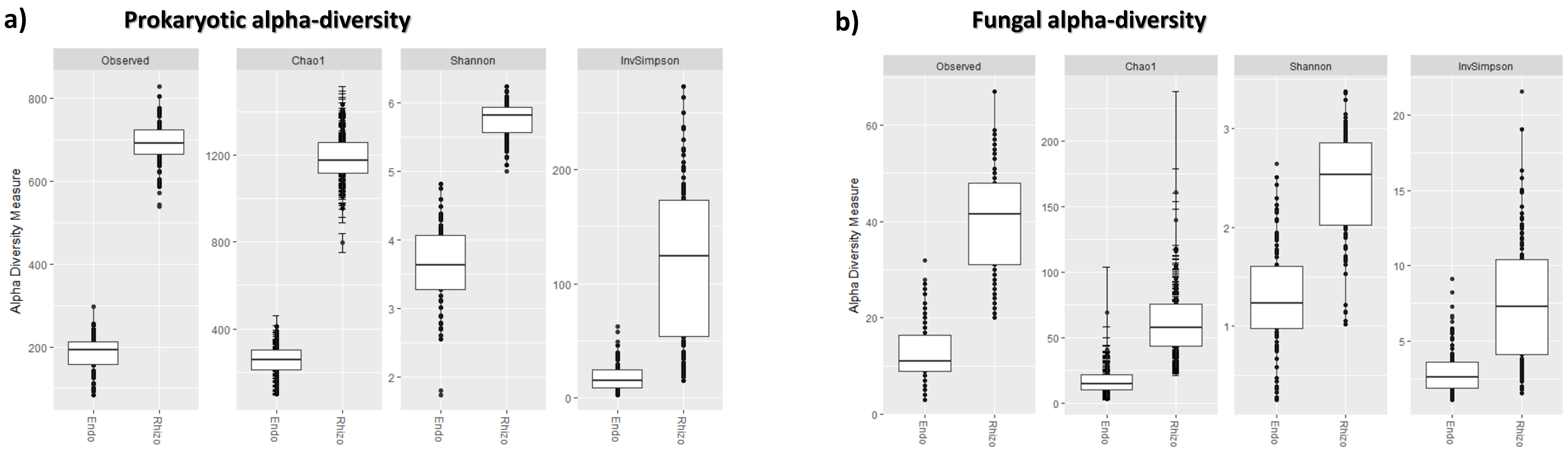

Figure S2. Microbial (bacterial a and b; fungal c and d) normalized alpha diversity indices of each sample clustered by cultivars in both compartments (a and c endosphere; b and d rhizosphere).

### a) Bacterial Endosphere

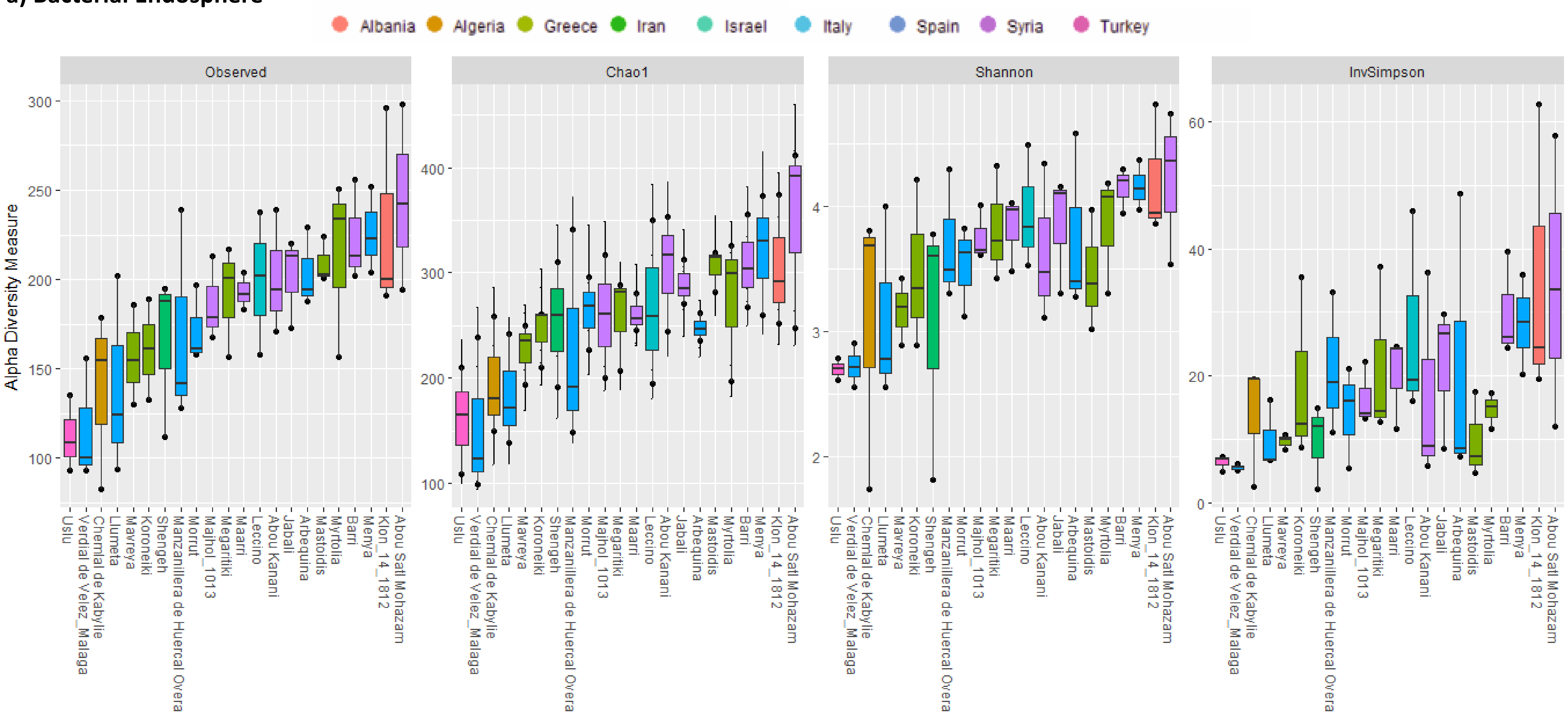

Figure S2. Microbial (bacterial a and b; fungal c and d) normalized alpha diversity indices of each sample clustered by cultivars in both compartments (a and c endosphere; b and d rhizosphere).

**b) Bacterial Rhizosphere**

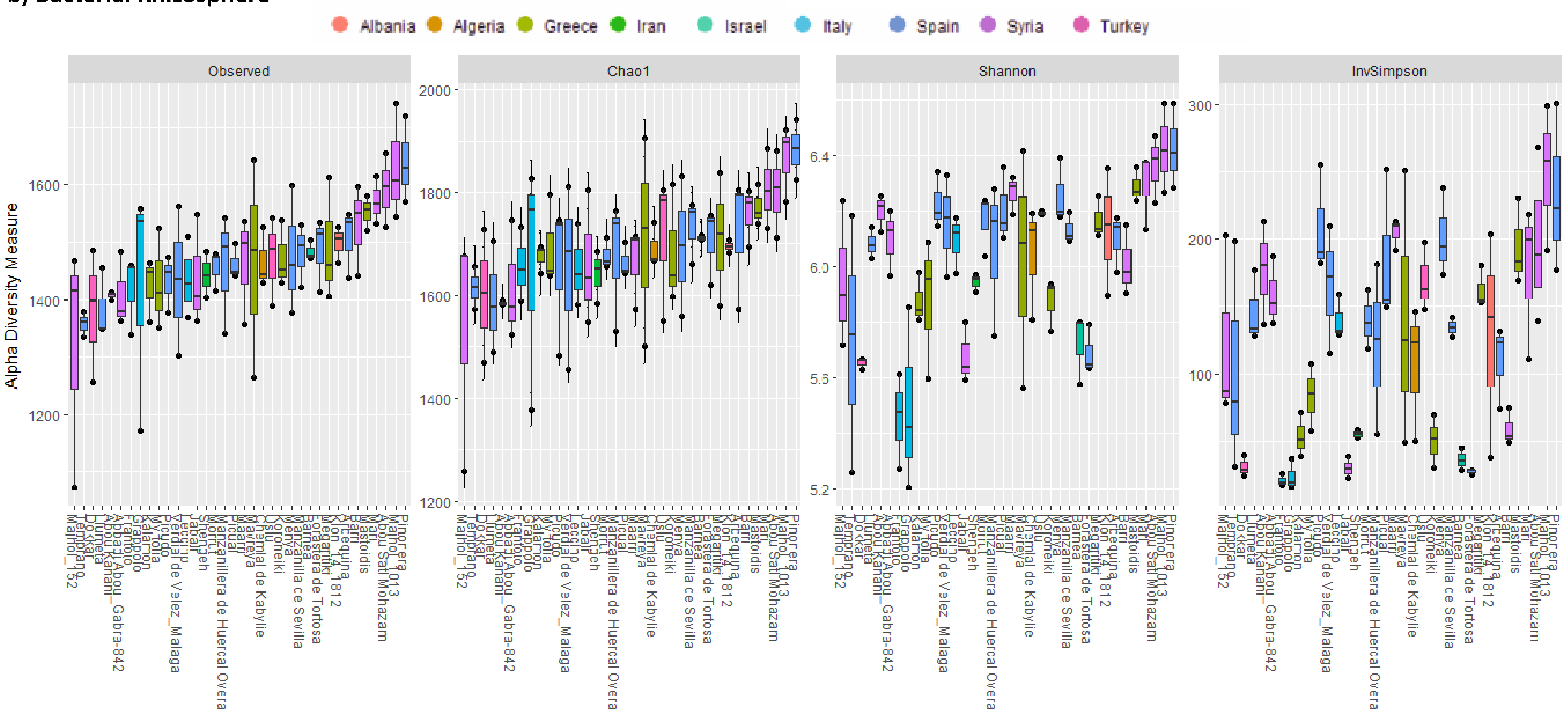

Figure S2. Microbial (bacterial a and b; fungal c and d) normalized alpha diversity indices of each sample clustered by cultivars in both compartments (a and c endosphere; b and d rhizosphere).

### c) Fungal Endosphere

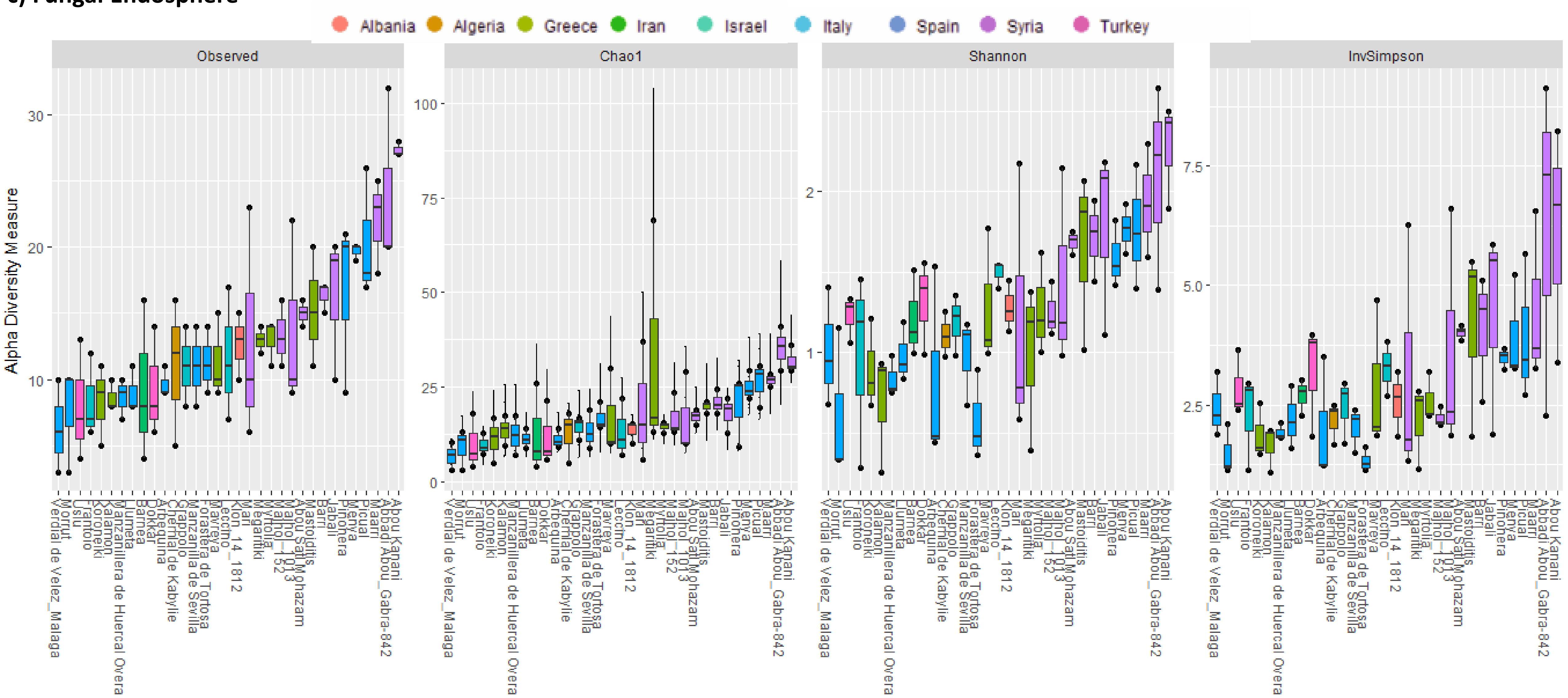

Figure S2. Microbial (bacterial a and b; fungal c and d) normalized alpha diversity indices of each sample clustered by cultivars in both compartments (a and c endosphere; b and d rhizosphere).

**d) Fungal Rhizosphere**

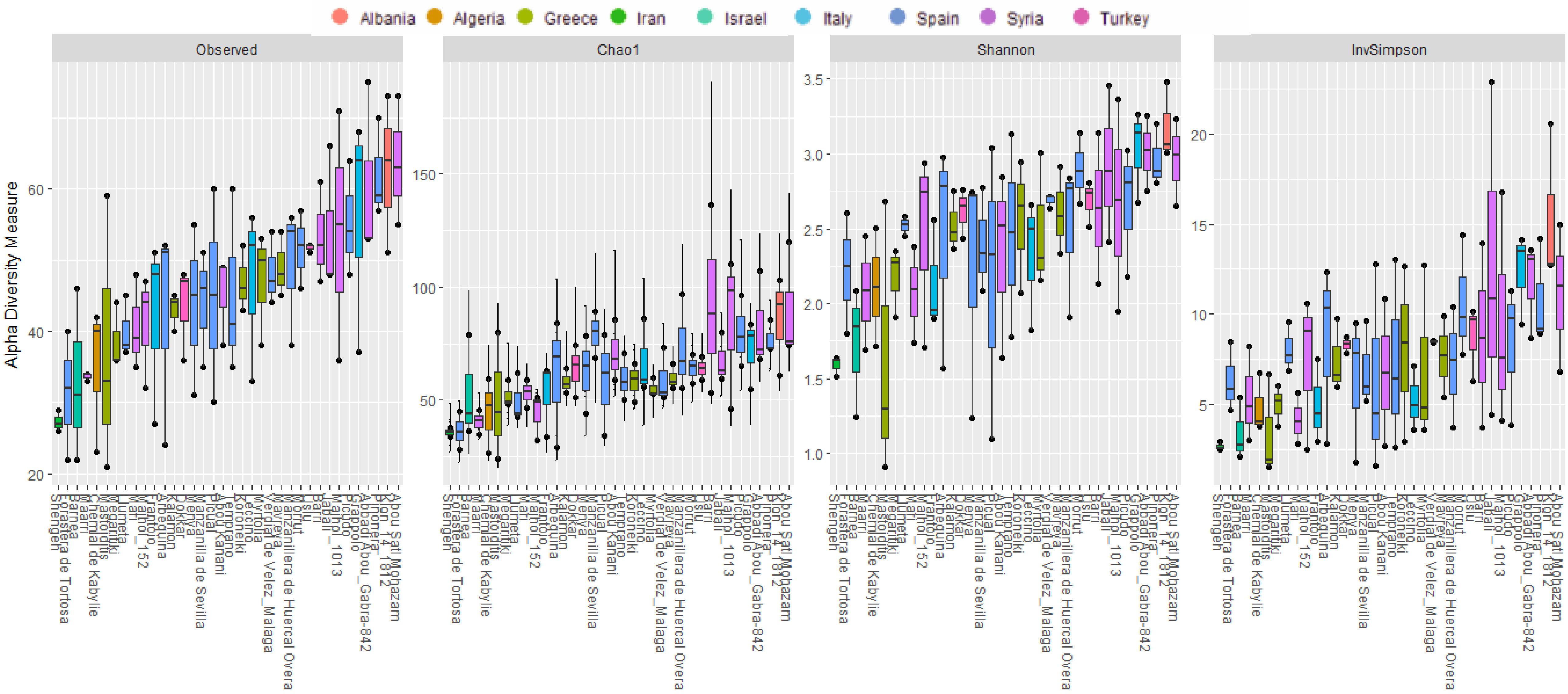

Figure S3. Statistically significant endophytic bacterial genera by cultivar (a, b) and the main bacterial genera in the endosphere (c).

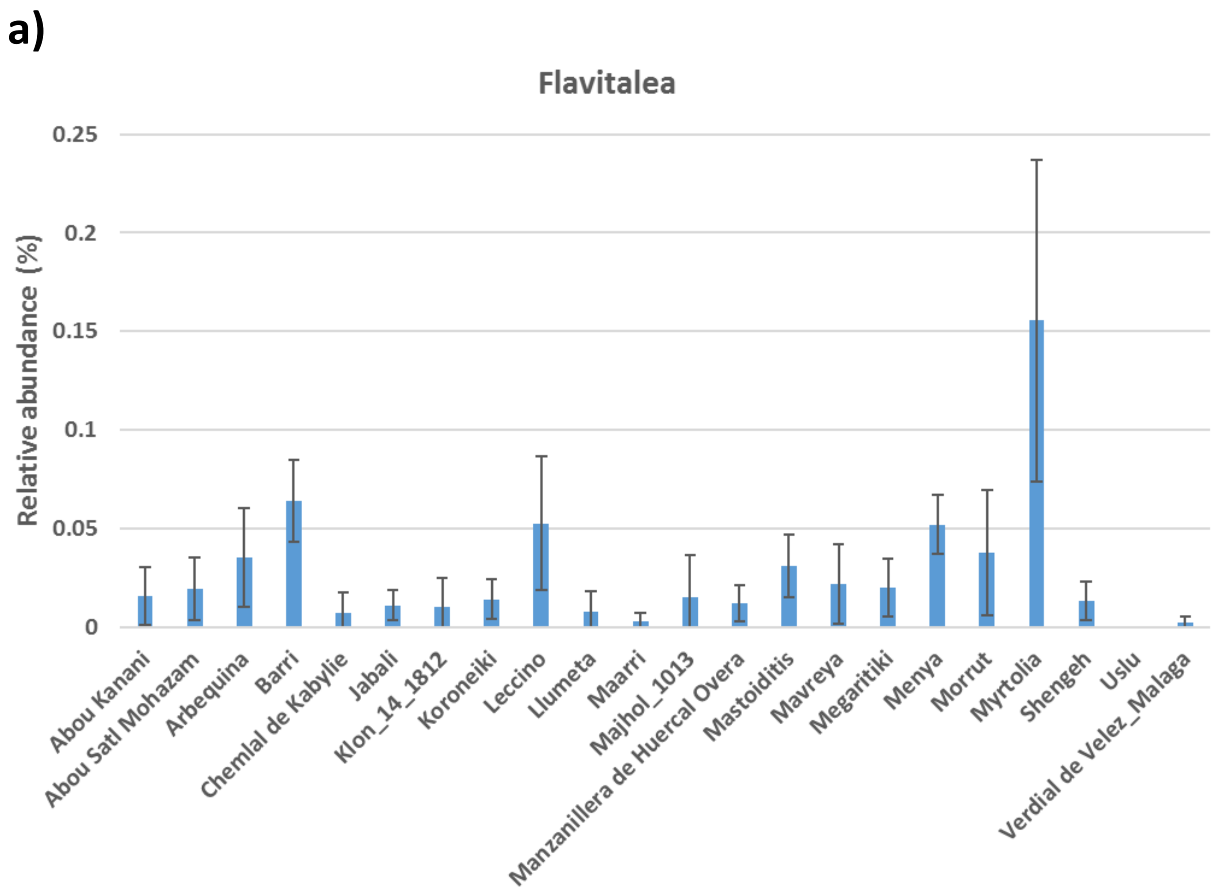


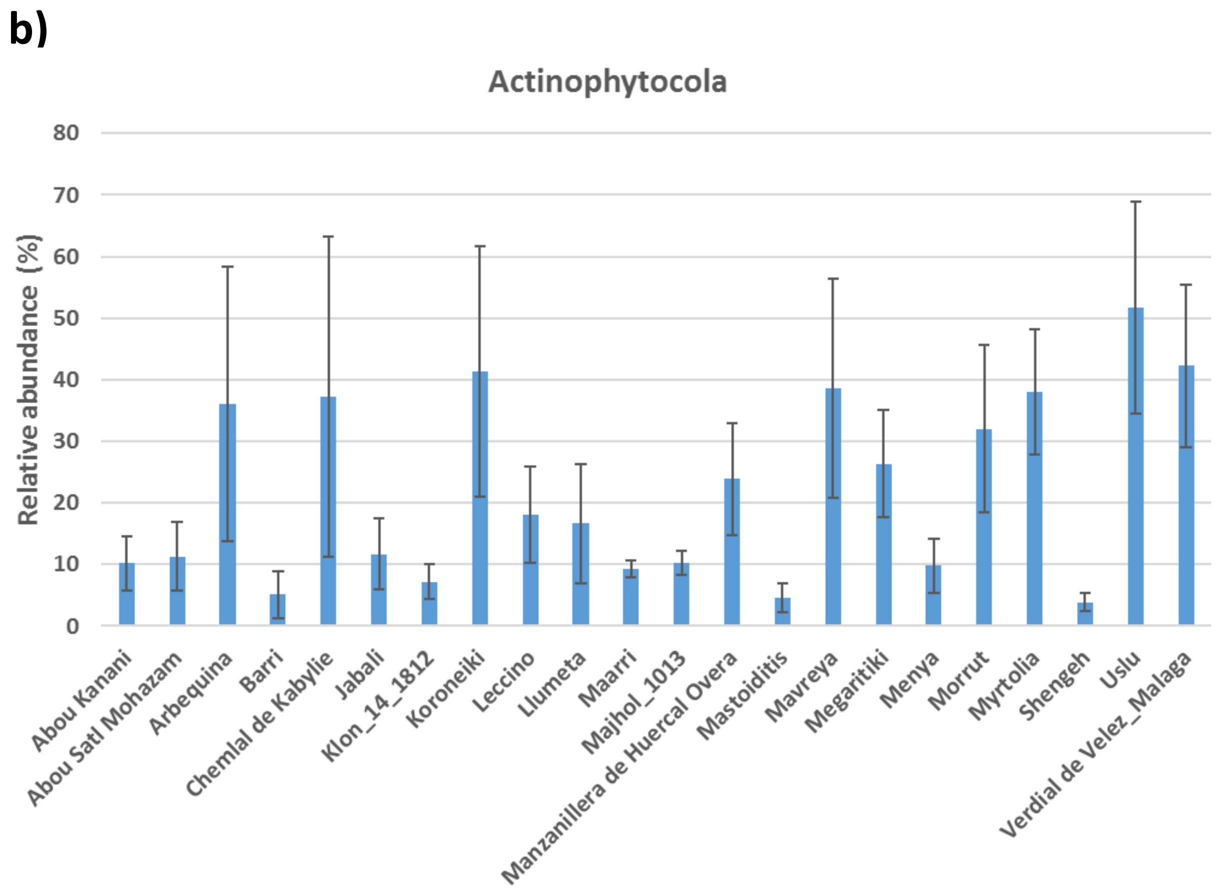

Figure S3. Statistically significant endophytic bacterial genera by cultivar (a, b) and the main bacterial genera in the endosphere (c).

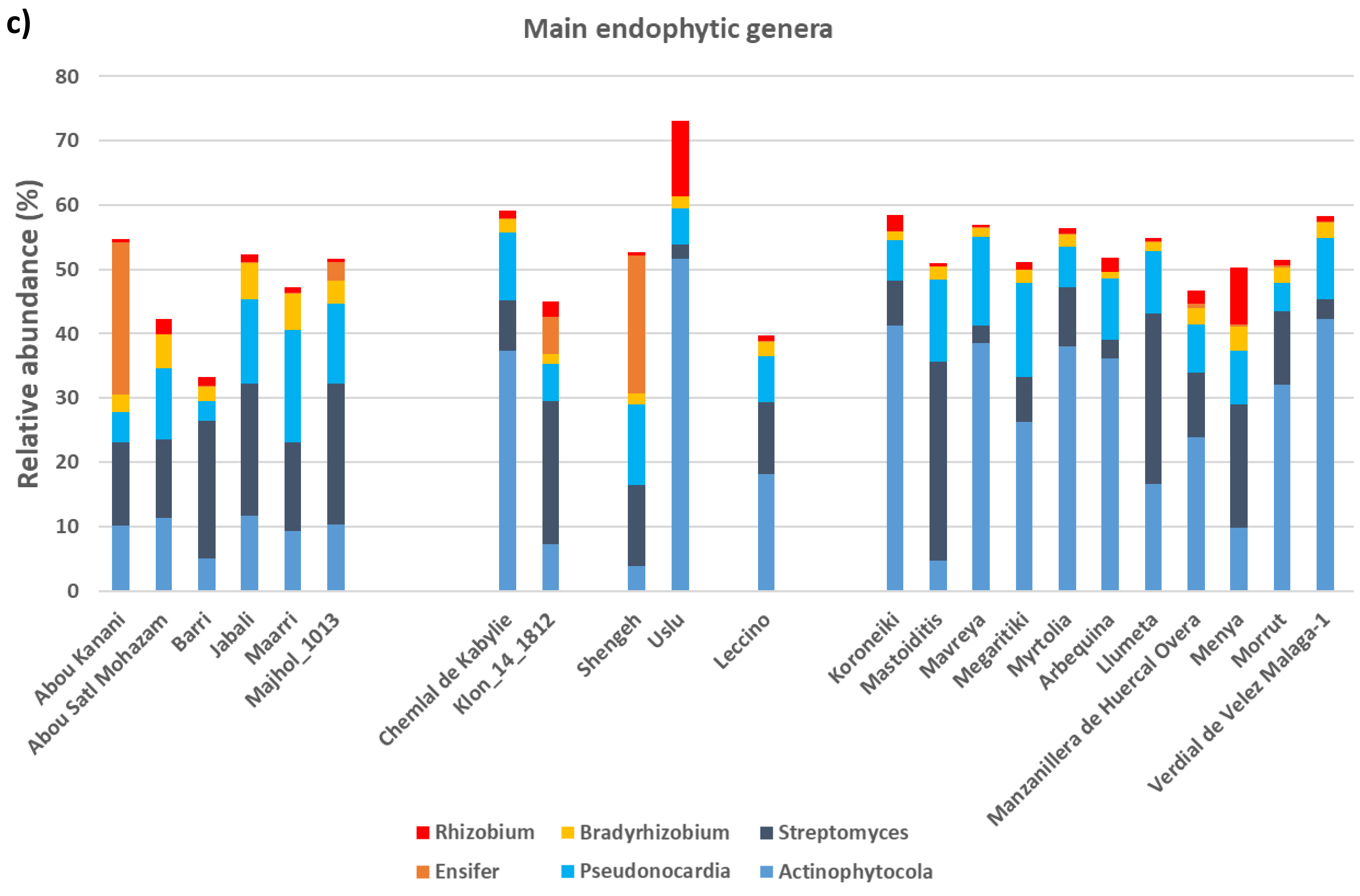

Figure S4. Main bacterial genera in the rhizosphere.

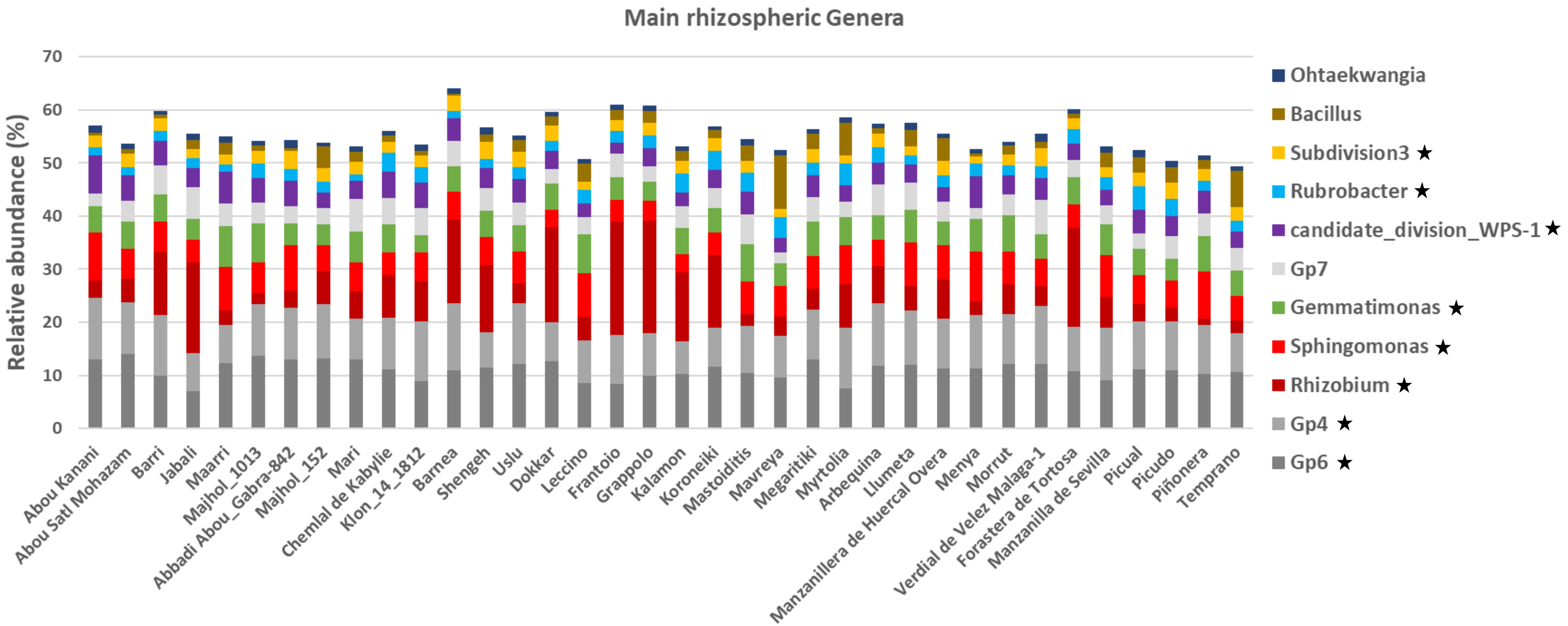

Figure S5. Statistically significant fungal endophytic (a) and rhizosphere (b) genera by cultivar and the main fungal genera in the endosphere (c) and the rhizosphere (d).

a)

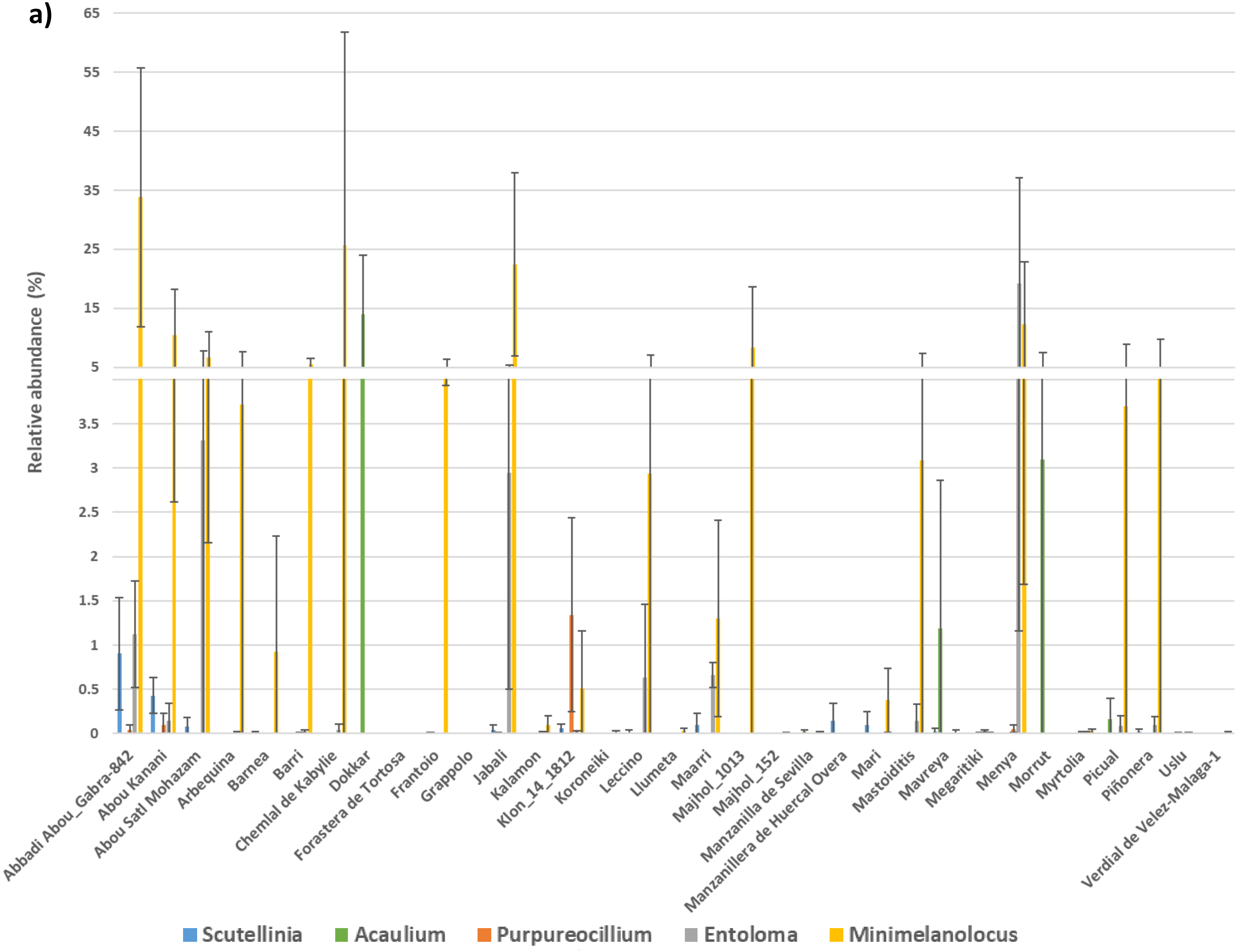

Figure S5. Statistically significant fungal endophytic (a) and rhizosphere (b) genera by cultivar and the main fungal genera in the endosphere (c) and the rhizosphere (d).

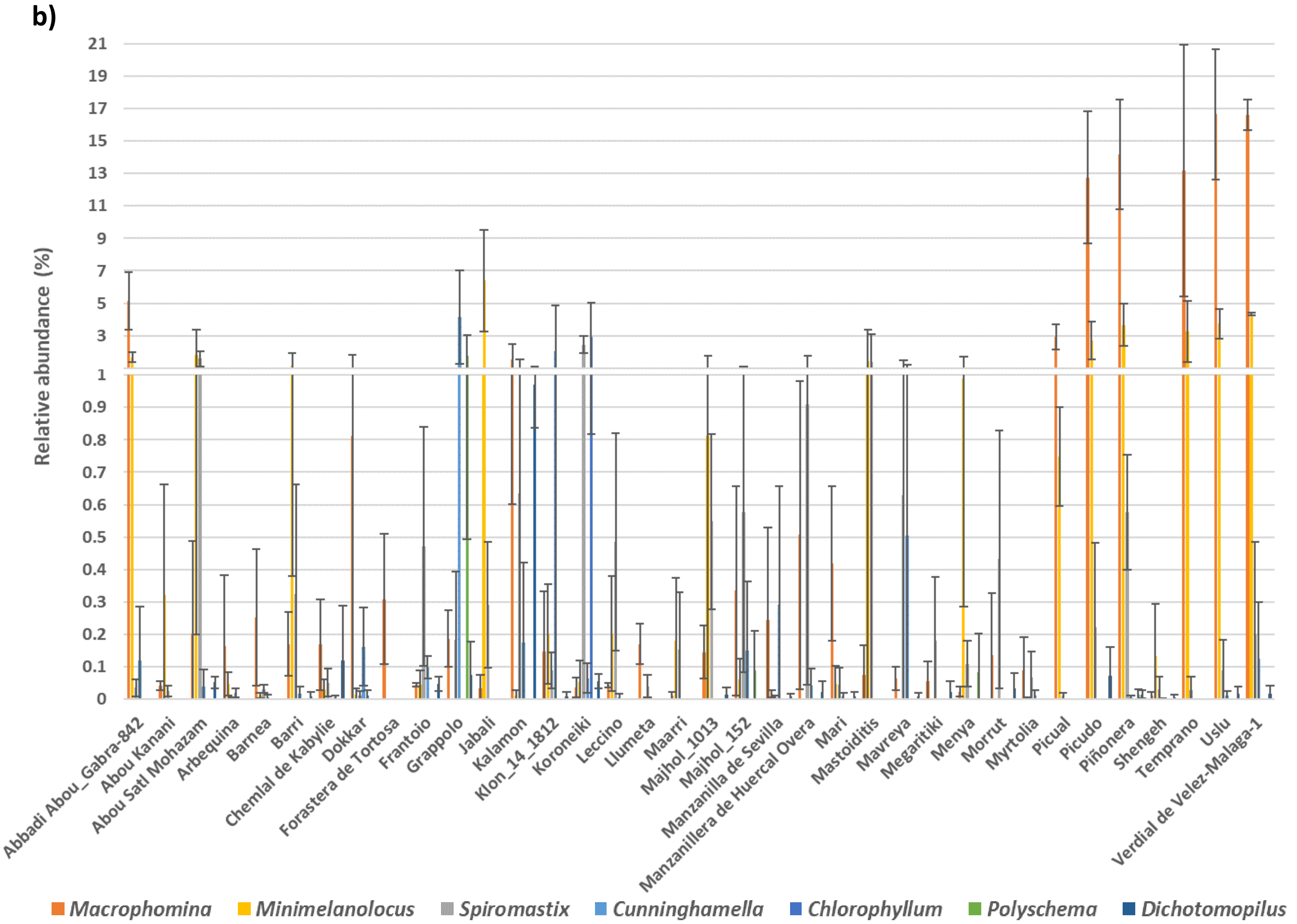

Figure S5. Statistically significant fungal endophytic (a) and rhizosphere (b) genera by cultivar and the main fungal genera in the endosphere (c) and the rhizosphere (d).

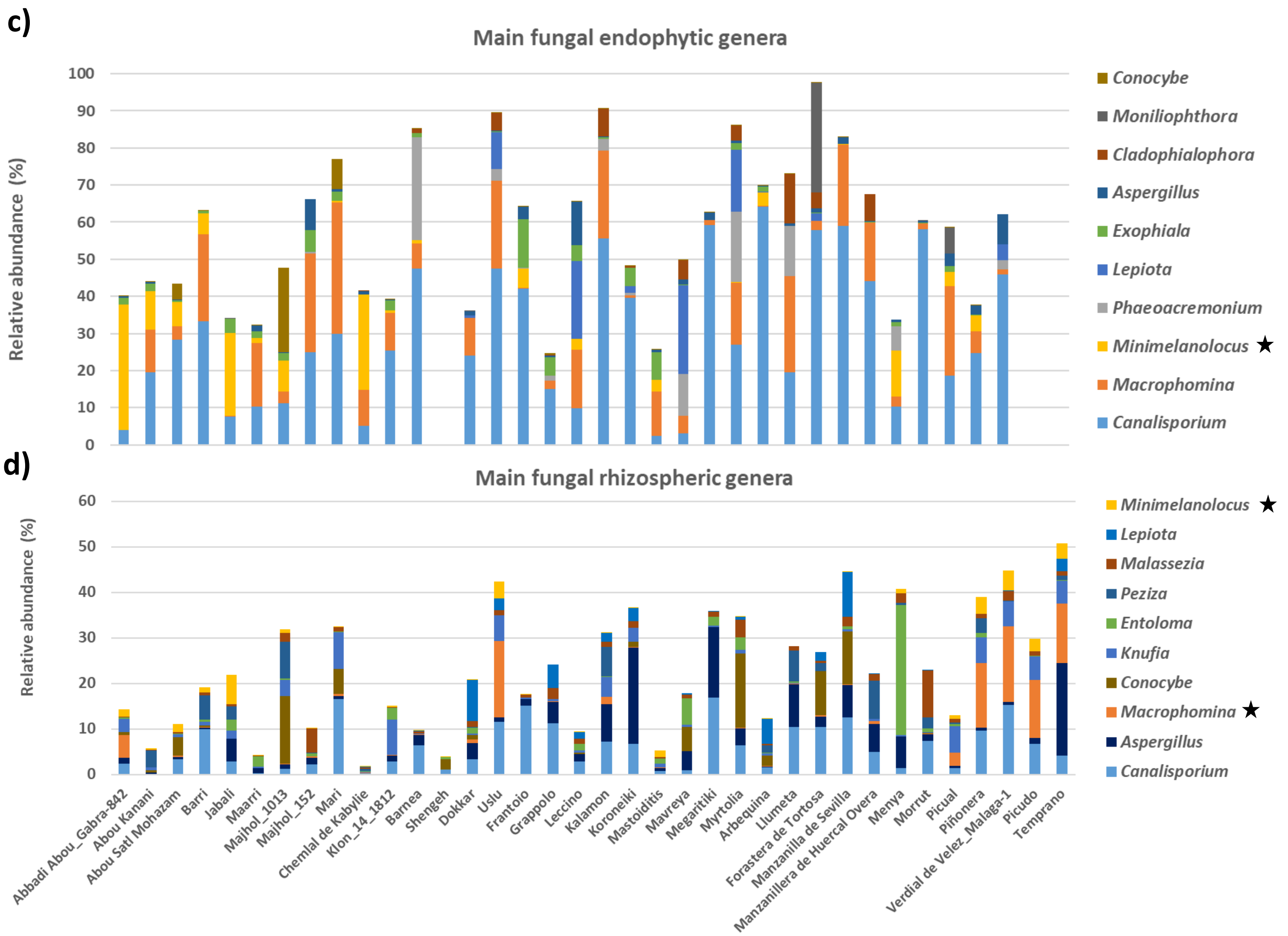

Table S5. Physicochemical properties of the soil from the World Olive Germplasm Collection (Córdoba, Spain)

| **Parameter** | **Mean values[1]** |
|---|---|
| Cation Exchange Capacity (meq /100 g) | 13.97 ± 4.27 |
| Calcium (mEq /100 g) | 9.42 ± 4.10 |
| Magnesium (mEq /100 g) | 3.33 ± 1.14 |
| Sodium (mEq /100 g) | 0.32 ± 0.05 |
| Potassium (mEq /100 g) | 0.90 ± 0.24 |
| Carbonates (%) | 15.93 ± 6.67 |
| Active lime (%) | 2.06 ± 1.09 |
| Assimilable phosphorus (p.p.m.) | 26.27 ± 10.37 |
| Organic matter (%) | 1.05 ± 0.53 |
| Organic nitrogen (%) | 0.09 ± 0.04 |
| pH 1 / 2.5 | 8.43 ± 0.19 |
| pH (in KCl) | 7.54 ± 0.14 |
| Exchangeable potassium (p.p.m.) | 380.5 ± 101.7 |
| Clay (%) | 22.16 ± 5.60 |
| Sand (%) | 40.97 ± 7.86 |
| Silt (%) | 36.87 ± 3.53 |

[1]Based on 10 samples (± standard deviation)
